# The origin of floral quartet formation - Ancient exon duplications shaped the evolution of MIKC-type MADS-domain transcription factor interactions

**DOI:** 10.1101/2022.12.23.521771

**Authors:** Florian Rümpler, Chiara Tessari, Lydia Gramzow, Christian Gafert, Marcus Blohs, Günter Theißen

## Abstract

During development of flowering plants, some MIKC-type MADS-domain transcription factors (MTFs) exert their regulatory function as heterotetrameric complexes bound to two sites on the DNA of target genes. This way they constitute „floral quartets“ or related „floral quartet-like complexes“ (FQCs), involving a unique multimeric system of paralogous protein interactions. Tetramerisation of MTFs is brought about mainly by interactions of keratin-like (K) domains. The K-domain associated with the more ancient DNA-binding MADS-domain during evolution in the stem group of extant streptophytes (charophyte green algae + land plants). However, whether this was sufficient for MTF tetramerisation and FQC formation to occur, remains unknown. Here, we provide biophysical and bioinformatic data indicating that the ancestral MTFs were not able to form FQCs. According to our data, FQC formation originated in the stem group of land plants in a sublineage of MIKC-type genes termed MIKC^C^-type genes. In the stem group of this gene lineage, the duplication of the most downstream exon encoding the K-domain led to a C-terminal elongation of the second K-domain helix, thus generating the tetramerisation interface found in extant MIKC^C^-type proteins. In the stem group of the sister lineage of the MIKC^C^-type genes, termed MIKC*-type genes, the duplication of two other exons of the K-domain occurred, extending the K-domain at its N-terminal end. Our data indicate that this structural change prevents heterodimerisation between MIKC^C^-type and MIKC*-type proteins. This way, two largely independent gene regulatory networks could be established, featuring MIKC^C^-type or MIKC*-type proteins, respectively, that control different aspects of plant development.

## INTRODUCTION

MADS-box genes, encoding MADS-domain transcription factors (MADS-TFs), constitute a conserved family of developmental control genes that are found in almost all eukaryotic organisms and that fulfil important functions in animals, plants and fungi. MADS-TFs of animals, fungi and some MADS-TFs of plants have in common only the presence of the conserved DNA-binding MADS-domain that specifically binds to cis-regulatory DNA elements termed CArG-box (for ‘C-A-rich-G’). MIKC-type MADS-TFs (termed MTFs henceforth for simplicity) of plants in addition carry a conserved keratin-like protein-protein interaction domain giving rise to the eponymous ‘MIKC’ domain architecture comprising an N-terminal MADS-domain (M) followed by an intervening- (I), keratin-like (K), and C-terminal (C) domain (Kaufmann, et al. 2005; Theißen and Gramzow 2016). Exactly when and how the K-domain originated is not known yet. However, as (almost) all streptophytes (that is charophyte green algae + land plants) carry MIKC-type genes, whereas chlorophytes (another group of green algae and sister group of streptophytes) do not, it appears likely that the K-domain evolved in the stem group of extant streptophytes (Tanabe, et al. 2005; Derelle, et al. 2006). In land plants MTFs subdivide into the two subfamilies of MIKC^C^- and MIKC*-type proteins that display structural differences within the I- and K-domain (Henschel, et al. 2002; Kaufmann, et al. 2005) and that have been shown to preferentially bind to different types of CArG-boxes termed SRF-type (consensus sequence 5’-CC(A/T)_6_GG-3’) and N10-type (5’-C(A/T)_8_G-3’), respectively (Verelst, et al. 2007; Zobell, et al. 2010; Wu, et al. 2011). Both land plant specific subfamilies most likely trace back to the duplication of an ancestral MIKC-type MADS-box gene prior to the transition of plants to land (Gramzow and Theißen 2010). However, the exact phylogenetic relationship of charophyte MIKC-type genes and MIKC^C^- and MIKC*-type genes of land plants as well as the structural features that distinguish both land plant subfamilies have been discussed controversially in the literature (Henschel, et al. 2002; Tanabe, et al. 2005; Kwantes, et al. 2012; Nishiyama, et al. 2018; Thangavel and Nayar 2018). Whereas charophytes usually carry only one MIKC-type gene, their number heavily increased during land plant evolution giving rise to 39 MIKC^C^- and 6 MIKC*-type genes in *Arabidopsis thaliana*, for example (Gramzow and Theißen 2010).

In seed plants, MTFs are involved in the control of a plethora of developmental processes, ranging from root development to floral induction, flower and fruit development of angiosperms (Smaczniak, Immink, Angenent, et al. 2012). Thereby, the transcription factors do not bind to DNA alone but as homo- or heteromeric complexes of two or four proteins. According to large-scale interaction-data, MIKC^C^- and MIKC*-type proteins form two mostly separated protein-protein interaction networks that predominantly control aspects of sporophyte (MIKC^C^) and gametophyte (MIKC*) development, respectively (de Folter, et al. 2005; Immink, et al. 2009; Smaczniak, Immink, Angenent, et al. 2012). The best studied complexes of MTFs are part of so called ‘floral quartets’ that are presumed to define the identities of the different floral organs during flower development of angiosperms (Theißen 2001; Theißen and Saedler 2001; Theißen, et al. 2016). A floral quartet consists of a tetramer of four MTFs simultaneously bound to two separated DNA-binding sites via looping the DNA in between. The simultaneous binding to both binding sites and the DNA-loop formation thereby presumably activate target gene expression by yet largely hypothetical epigenetic mechanisms (Mendes, et al. 2013; Theißen, et al. 2016). The existence of MTF tetramers has been shown not to be limited to floral homeotic proteins controlling flower development (Wang, et al. 2010; Ruelens, et al. 2017). Rather it appears likely that at least most MIKC^C^-type proteins of seed plants can be incorporated into floral quartet like complexes (FQCs) (Espinosa-Soto, et al. 2014; Puranik, et al. 2014; Rümpler, et al. 2018) and it has been hypothesized that the ability of MIKC^C^-type proteins to tetramerise was probably an important precondition for establishing and diversifying sharp developmental switches (Theißen, et al. 2016).

The protein-protein interactions that facilitate MIKC^C^-type protein dimerization and tetramerization are mainly mediated by the K-domain (Yang and Jack 2004; Melzer, et al. 2009; Puranik, et al. 2014; Rümpler, et al. 2018). According to structural data of the floral homeotic MIKC^C^-type protein SEPALLATA 3 (SEP3) from *A. thaliana*, the K-domain folds into two amphipathic α-helices that constitute coiled-coils (Puranik, et al. 2014), a common and well-studied class of protein-protein interaction domains (Lupas and Gruber 2005). The first helix and the N-terminal part of helix 2 strengthen the dimeric interaction between two SEP3 monomers bound to one DNA-binding site. The C-terminal part of helix 2 allows the interaction of two DNA-bound dimers thus facilitating FQC formation (Puranik, et al. 2014; Rümpler, et al. 2018). Due to its high sequence conservation, it is presumed that the K-domains of most MIKC^C^-type proteins follow a structure that is very similar to that determined for SEP3 (Rümpler, et al. 2018).

There is growing evidence that FQC formation is widespread among seed plant MIKC^C^-type proteins (Espinosa-Soto, et al. 2014; Rümpler, et al. 2018) and that tetramerization is of high functional relevance (Mendes, et al. 2013; Hugouvieux, et al. 2018). However, neither is anything known about FQC formation capabilities of MIKC*-type proteins nor when during evolution MTFs ‘learned’ to constitute quartets and which evolutionary changes facilitated this important ability. Here, we present bioinformatical and molecular biophysical data which demonstrate that FQC origin most likely coincides with the transition of plants to land more than 500 million years ago. FQC formation was likely facilitated by the duplication of a K-domain exon of an ancestral MIKC^C^-type gene. In contrast, two other K-domain exons very likely got duplicated during early MIKC*-type gene evolution, resulting in structural differences in the protein-protein interaction interfaces of MIKC^C^- and MIKC*-type proteins eventually preventing heteromeric interactions between members of both subfamilies. We hypothesize that the ancient exon duplications created the molecular prerequisites for the evolution of effective and diverse developmental switches and the establishment of two independent interaction networks controlling sporophyte and gametophyte development of land plants, respectively.

## RESULTS

### MIKC^C^-type protein representatives from ferns, lycophytes and mosses form FQCs

The ability of MTFs to constitute FQCs has so far only been tested for a limited set of MIKC^C^-type proteins from seed plants (Melzer and Theißen 2009; Melzer, et al. 2009; Wang, et al. 2010; Smaczniak, Immink, Muino, et al. 2012; Jetha, et al. 2014; Ruelens, et al. 2017; Rümpler, et al. 2018). To investigate the tetramerisation capabilities and DNA-binding of representative MIKC^C^-type proteins from ferns, lycophytes and bryophytes we used a well-established electrophoretic mobility shift assay (EMSA) (Melzer and Theißen 2009; Melzer, et al. 2009; Rümpler, et al. 2018). The rationale of our approach is described in detail in Material and Methods. We synthesized and cloned the coding sequences of *PHYSCOMITRELLA PATENS MADS 1* (*PPM1*) from the moss *Physcomitrium patens* (formerly *Physcomitrella patens*), *SELAGINELLA MOELLENDORFFII MADS 3* (*SmMADS3*) from the lycophyte *Selaginella moellendorffii*, and *CERATOPTERIS RICHARDII MADS 3* (*CRM3*) from the fern *Ceratopteris richardii*. The encoded proteins were produced *in vitro* and their ability to form FQCs was analysed via EMSA using a radioactively labelled DNA probe that carried two SRF-type CArG-boxes of the sequence 5’-CCAAATAAGG-3’ in a distance of 63 bp (about six helical turns) (Melzer and Theißen 2009; Melzer, et al. 2009; Rümpler, et al. 2018). When increasing amounts of *in vitro* translated protein were co-incubated with a constant amount of DNA, a single retarded fraction of reduced electrophoretic mobility was observed for all of the investigated MIKC^C^-type proteins (Figure 1a,c,e and Supplementary Figure 1a,c,e). This is in contrast to most seed plant MIKC^C^-type proteins investigated so far, which usually produce two retarded fractions (at least for low amounts of applied protein), which constitute complexes of two and four proteins bound to DNA, respectively (Melzer and Theißen 2009; Melzer, et al. 2009; Wang, et al. 2010; Jetha, et al. 2014; Rümpler, et al. 2018).

**Figure 1:**
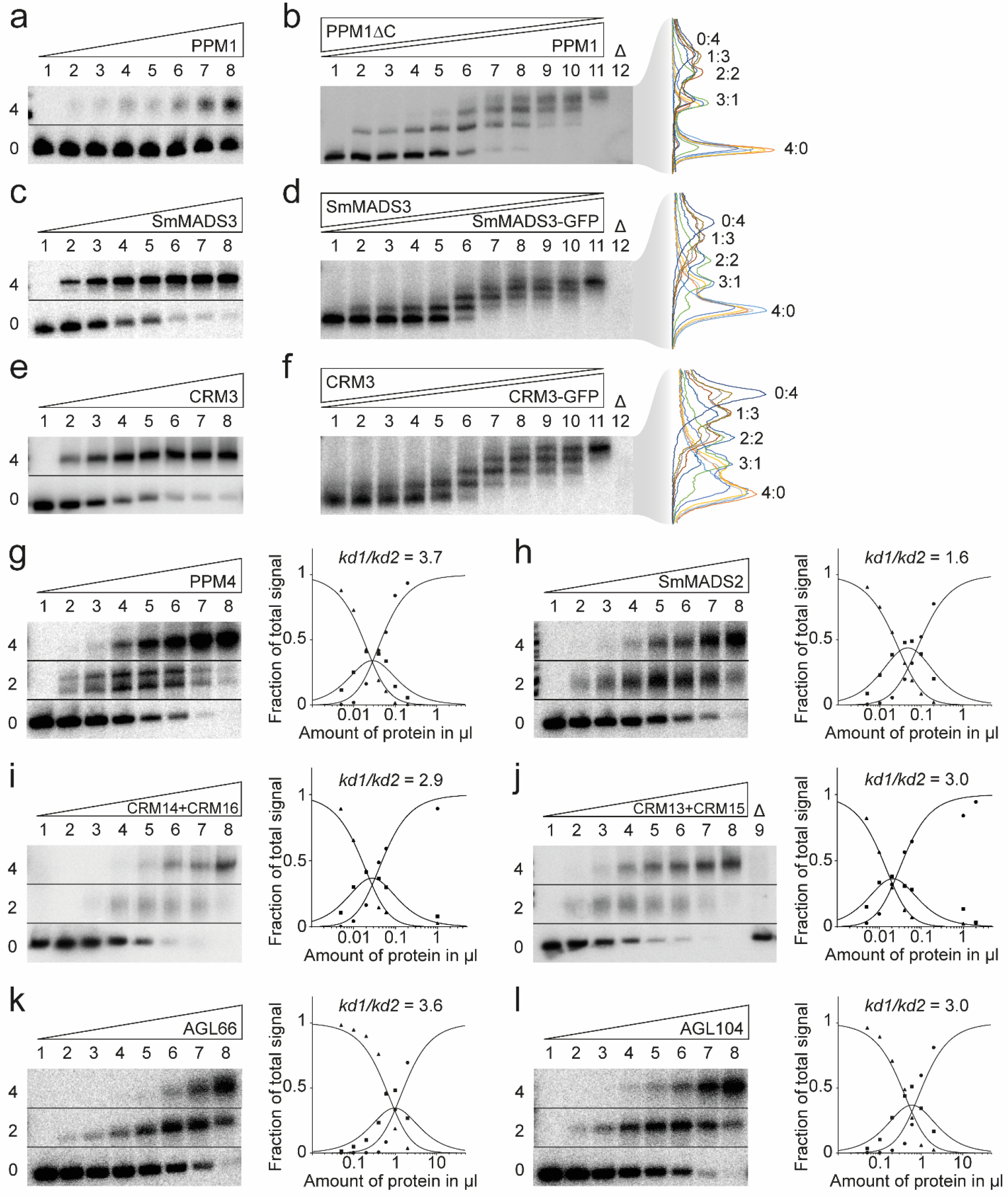
FQC formation capabilities of MIKC^C^- and MIKC*-type MADS-TFs from *P. patens, S. moellendorffìi, C. richardii* and *A. thaliana*. (a, c, e) Increasing amounts of *in vitro* transcribed/translated (a) PPM1, (c) SmMADS3 and (e) CRM3 protein, respectively, were co-incubated together with constant amounts of radioactively labelled DNA probe carrying two SRF-type CArG-boxes. Two fractions of different electrophoretic mobility occur – a fraction of high electrophoretic mobility, constituting unbound DNA probe (labelled with ‘0’), and a retarded fraction constituting a DNA probe bound by four proteins (‘4’). (b, d, f) To determine the stoichiometry of the protein-DNA complexes observed in a, c, and e, (b) PPM1, (d) SmMADS3, and (f) CRM3 wild type proteins were co-expressed at different ratios with PPM1ΔC, SmMADS3-GFP, and CRM3-GFP, respectively, and co-incubated together with constant amounts of the same radioactively labelled DNA probe as in a, c, and e. For all proteins five retarded fractions of different electrophoretic mobility occurred. An overly of measured signal intensities of the individual lanes is shown on the right. Each peak of the graph is labelled according to the ratio of wild type and truncated/elongated protein of the corresponding fraction. (g-l) Increasing amounts of *in vitro* transcribed/translated (g) PPM4, (h) SmMADS2, (i) CRM14 + CRM16, (j) CRM13 + CRM15, (k) AGL66, and (l) AGL104 was co-incubated together with constant amounts of radioactively labelled DNA probe carrying two N10-type CArG-boxes. Three fractions of different electrophoretic mobility occur – a fraction of high electrophoretic mobility constituting unbound DNA probe (labelled with ‘0’), a fraction of intermediate electrophoretic mobility constituting a DNA probe bound by a single protein dimer (‘2’) and a fraction of low electrophoretic mobility constituting a DNA probe bound by four proteins (‘4’). Signal intensities of different fractions were measured and plotted against the amount of applied protein (triangles, free DNA; squares, DNA probe bound by two proteins; circles, DNA probe bound by four proteins). Graphs were fitted according to equations (1) to (3) described in Materials and Methods to eventually quantify and express FQC formation capabilities by *k_d1_/k_d2_*. In case of (g) PPM4, a double band was observed for the fraction of intermediate electrophoretic mobility, likely caused by different conformations of the DNA-bound protein dimer. As negative control 2 μl of *in vitro* transcription/translation mixture loaded with the empty pTNT plasmid were added to the binding reaction (‘Δ’). (a, c, e, g-l) Applied amounts of *in vitro* transcription/translation products were (lane 1 to 8) 0, 0.05, 0.1, 0.2, 0.4, 0.6, 1, and 2 μl, whereby CRM3, PPM4, SmMADS2, CRM14+CRM16, and CRM13+CRM15 were prediluted 1:10 and PPM1 and SmMADS3 were prediluted 1:20 with BSA (10 mg ml^-1^). (b, d, f) 3 μl of *in vitro* transcription/translation product were applied to each lane. Ratios of both template plasmids used for *in vitro* transcription/translation were (lane 1 to 11): 0:1, 1:9, 1:7, 1:5, 1:3, 1:1, 3:1, 5:1, 7:1, 9:1, 1:0.

The electrophoretic mobility of the single retarded fraction observed for the fern, lycophyte and moss MIKC^C^-type proteins suggests a complex of four DNA-bound proteins. However, to determine the stoichiometry of the observed complexes more accurately, as had been previously established (Melzer et al., 2009), we generated a truncated version of PPM1 (PPM1ΔC) that only comprises MADS-, I- and K-domain of PPM1 but lacks most parts of the C-terminal domain (amino acids 175-283) and co-incubated variable amounts of wild type and C-terminally truncated protein together with labelled DNA probe. Following the assumption that all PPM1 full-length proteins within the DNA-bound complex can be substituted by a truncated PPM1ΔC protein, the number of fractions with different electrophoretic mobility reveals the stoichiometry of the DNA-bound complex. Similar to PPM1 full length protein, PPM1ΔC alone produced a single retarded fraction but with a higher electrophoretic mobility due to the reduced protein size (Figure 1b, lane 1 and Supplementary Figure 1b). If PPM1 and PPM1ΔC were mixed at different ratios, in total five retarded fractions of different electrophoretic mobility occurred (Figure 1b, lanes 2-10 and Supplementary Figure 1b), representing DNA-bound complexes of PPM1/PPM1ΔC in protein ratios of 0:4, 1:3, 2:2, 3:1 and 4:0, respectively. This indicates that the single retarded fraction observed in Figure 1a constitutes a DNA probe bound by four PPM1 proteins. Since in Figure 1a no fraction of intermediate electrophoretic mobility (i.e. a single protein dimer bound to DNA) was observed even at low amounts of applied protein, PPM1 seems to form FQCs with very high affinity under our experimental conditions (i.e. it almost immediately occupies both DNA-binding sites), either because dimerization of DNA-bound MTF dimers occurs in a highly cooperative way, or even MTF tetramers are formed in free solution before binding to DNA.

Because SmMADS3 and CRM3 have considerably shorter C-terminal domains than PPM1, we conducted the stoichiometry tests with elongated instead of shortened protein versions. We generated elongated versions of SmMADS3 and CRM3 via fusion to the green fluorescent protein (SmMADS3-GFP and CRM3-GFP) and co-incubated variable amounts of wild type and GFP-fused proteins together with labelled DNA probe. Similar to PPM1, we observed in total five retarded fractions of different electrophoretic mobility for SmMADS3/SmMADS3-GFP and CRM3/CRM3-GFP, respectively (Figure 1d,f and Supplementary Figure 1d,f), demonstrating that also SmMADS3 and CRM3 bind to DNA immediately as tetramer and thus form FQCs with very high affinity under the applied conditions.

### All investigated MIKC*-type proteins are unable to form FQCs

To examine the FQC formation capabilities of MIKC*-type proteins, we cloned the coding sequences of *PPM4* from *P. patens, SmMADS2* from *S. moellendorffii, CRM13, CRM14, CRM15*, and *CRM16* from *C. richardii*, and *AGAMOUS-LIKE 66* (*AGL66*) and *AGL104* from *A. thaliana* and expressed the proteins *in vitro*. Because MIKC*-type proteins are known to preferentially bind to N10-type CArG-box sequences (Verelst, et al. 2007; Zobell, et al. 2010; Wu, et al. 2011), we tested all MIKC*-type proteins for their FQC formation capabilities against a radioactively labelled DNA-probe that carried two N10-type CArG boxes of the sequence 5’-CTATATATAG-3’ in a distance of 63 bp (about six helical turns). At low amounts of applied protein, all of the tested MIKC*-type proteins produced a fraction of intermediate electrophoretic mobility, and with increasing protein amounts an additional fraction of low electrophoretic mobility occurred (Figure 1g-l and Supplementary Figure 2). A similar DNA-binding behaviour is typically observed for seed plant MIKC^C^-type MADS-TFs (Melzer and Theißen 2009; Melzer, et al. 2009; Rümpler, et al. 2018), where the fraction of intermediate electrophoretic mobility corresponds to a DNA probe bound by two proteins (i.e. binding of a single dimer) and the fraction of low electrophoretic mobility constitutes a DNA probe bound by four proteins (i.e. binding of two dimers or one tetramer, respectively). By measuring the signal intensities of the three different fractions (free DNA, DNA probe bound by two proteins, and DNA probe bound by four proteins) the cooperative DNA-binding and thus FQC formation capabilities of the examined MIKC*-type proteins can be quantified and expressed as ration of the dissociation constants for binding of the first and the second dimer *k_d1_/k_d2_* (for details, see Materials and Methods), as described previously (Melzer, et al. 2009; Jetha, et al. 2014; Rümpler, et al. 2018). All investigated MIKC*-type proteins produced very low *k_d1_/k_d2_* values ranging from 1 to 7, indicating no or very weak positive interaction of two DNA-bound dimers and thus no ability to cooperatively form FQC under our experimental conditions. For comparison, similar tests with seed plant MIKC^C^-type proteins capable of forming FQCs, such as the SEP proteins (SEP1, SEP2, SEP3, SEP4) from *A. thaliana* or GNETUM GNEMON MADS 3 (GGM3), GGM9, GGM11 from *Gnetum gnemon*, resulted in cooperativity values of 100 or higher at similar experimental conditions indicating highly cooperative DNA-binding (Melzer, et al. 2009; Wang, et al. 2010; Jetha, et al. 2014).

### Duplications of distinct K-domain exons differentiate MIKC^C^- and MIKC*-type genes

MIKC^C^- and MIKC*-type proteins are known to differ with respect to their exon-intron structure, although there is no consensus in the literature as to whether these differences pertain to the I-domain and/or the K-domain (Henschel, et al. 2002; Tanabe, et al. 2005; Kwantes, et al. 2012; Nishiyama, et al. 2018; Thangavel and Nayar 2018). To investigate which sequence determinants might account for the observed differences in FQC formation capabilities, we conducted a large-scale exon homology analysis based on a multiple sequence alignment of all MIKC^C^- and MIKC*- type genes from *P. patens, S. moellendorffii, C. richardii*, and *A. thaliana*. With a few exceptions, all analysed MIKC^C^-type proteins are encoded by one MADS-domain exon, one I-domain exon, four K-domain exons and a variable number of C-terminal domain exons (Figure 2). K-domain exons were defined based on whether they were homologous to exons encoding for the K-domain of SEP3 from *A. thaliana* for which the crystal structure has been determined (Puranik, et al. 2014). In contrast to MIKC^C^-type proteins, the analysed MIKC*-type proteins are usually encoded by six instead of four K-domain exons. Exon homology analyses demonstrate that only three K-domain exons are shared by MIKC^C^- and MIKC*-type genes (highlighted by black arrows in Figure 2). The first three (most upstream) K-domain exons of MIKC*-type genes are not found among MIKC^C^-type genes and the last (most downstream) K-domain exon of MIKC^C^-type genes is not found among MIKC*-type genes (highlighted by red arrowheads in Figure 2).

**Figure 2:**
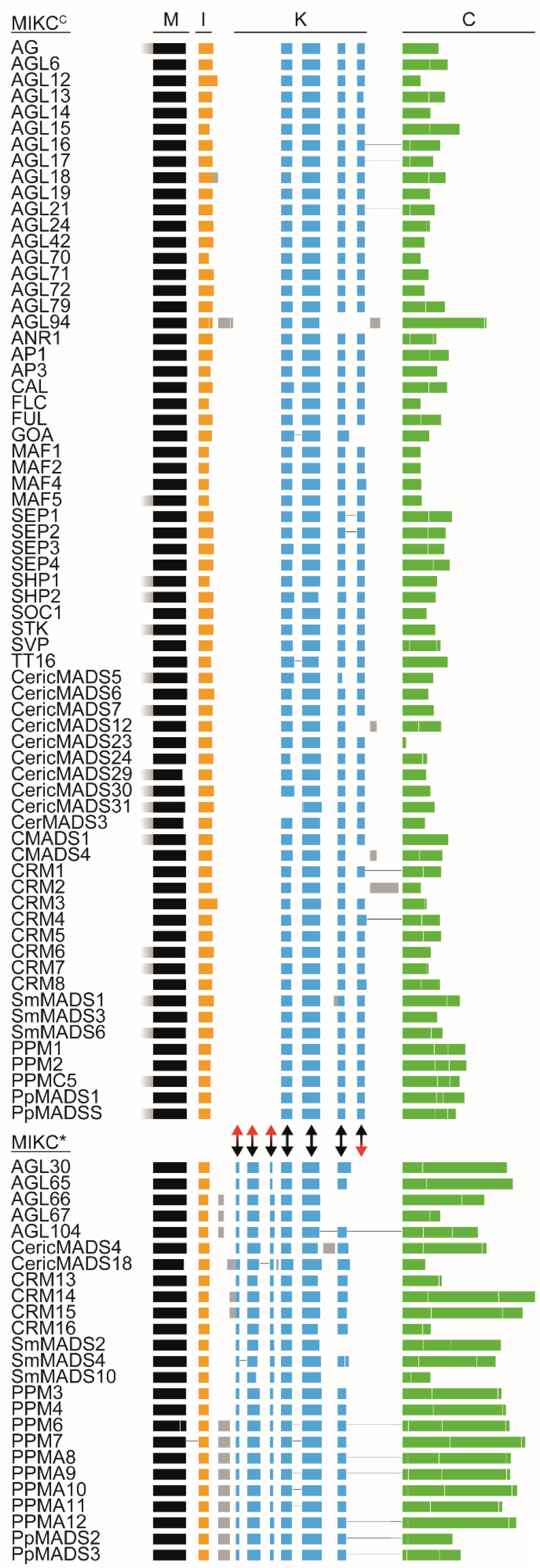
Exon homology of MIKC^C^- and MIKC*-type genes from *A. thaliana, C. richardii, S. moellendorffii* and *P. patens*. Coloured boxes represent coding exons of all MIKC^C^- and MIKC*-type genes from *A. thaliana, C. richardii, S. moellendorffii* and *P. patens*. Introns and non-coding exons are not shown. Exons encoding for the MADS-domain, the intervening domain (I-domain), the keratin-like domain (K-domain), and the C-terminal domain (C-domain) are shown in black, yellow, blue and green, respectively. Exons with uncertain homology assignment are colour-coded in grey. Exons encoding for MADS-, I, and K-domain were aligned according their homology based on a multiple sequence alignment of the encoded proteins back translated into a codon alignment (for details see Materials and Methods). Fused exons are connected by horizontal black lines. Two-headed arrows between MIKC^C^- and MIKC*-type genes illustrate presence (black) or absence (red arrowhead) of homologous exons in either of the two subfamilies.

The K-domain exons that differentiate MIKC^C^- and MIKC*-type genes show some remarkable peculiarities. The last two K-domain exons of MIKC^C^-type genes are of similar size, almost invariably encoding for 14 amino acids each. Analysis of the amino acid sequence encoded by these exons revealed that they also show similarity on the sequence level (Figure 3a). Likewise, the first four K-domain exons of MIKC*-type genes show similarities in size and sequence. The first and the third K-domain exons often encode for 7 amino acids each. The second and the fourth K-domain exons usually encode for 21 amino acids each (Figure 3b). Due to the similar size and the high level of sequence similarity, it appears very likely that the last two K-domain exons of MIKC^C^-type genes as well as the first four K-domain exons of MIKC*-type genes evolved by exon duplications of an ancestral and a pair of ancestral exons, respectively. As the hypothetically duplicated exons are present in almost all analysed MIKC^C^- and MIKC*-type genes, respectively, it appears likely that the exon duplication events took place in the stem lineage of MIKC^C^- and MIKC*-type genes, respectively and thus in the stem group of extant land plants.

**Figure 3:**
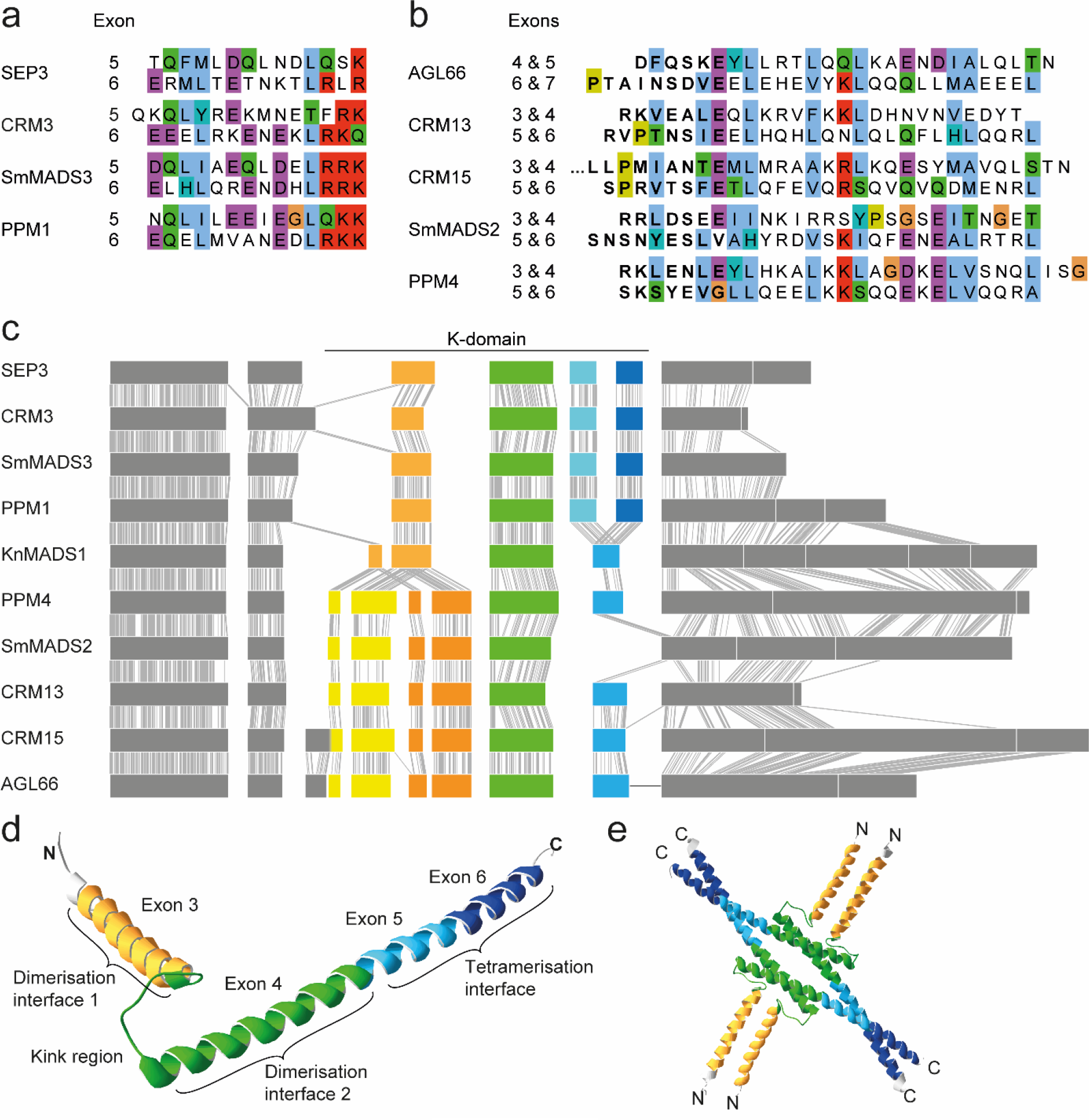
Similarity of K-domain exons hypothesized to be duplicated. (a, b) Multiple sequence alignment of the amino acids encoded by (a) exon 5 and 6 of the MIKC^C^-type genes *SEP3, CRM3, SmMADS3*, and *PPM1* and (b) exons 4 – 7 of the MIKC*-type gene *AGL66* and exons 3 – 6 of *CRM13, CRM15, SmMADS2*, and *PPM4*, respectively. (c) Exon-intron structure of the MIKC^C^- and MIKC*-type genes shown in panel a and b, respectively, together with the exon-intron structure of the charophyte MIKC-type gene *KnMADS1*. Homologous exons were aligned and identical nucleotide positions of neighbouring sequences are connected with solid grey lines to illustrate homology. K-domain exons shared by MIKC^C^-, MIKC*- and charophyte MIKC-type genes are colour-coded in green, hypothetically duplicated K-domain exons of MIKC^C^- and MIKC*-type genes are colour-coded in different shades of blue and yellow/orange, respectively. (d) X-ray crystal structure of the MIKC^C^-type protein SEP3 (PDB-ID: 4OX0). Subdomains encoded by exon 3, 4, 5, and 6 are colour-coded in yellow, green, light blue and dark blue, respectively. (e) Tetramer of four SEP3 K-domains following the same colour-coding as in d. Protein structure images were generated with Swiss-PdbViewer (Guex and Peitsch 1997).

### Charophyte MIKC-type genes show no duplications of K-domain exons

Land plant MIKC^C^- and MIKC*-type genes very likely evolved by a gene duplication of an ancestral MIKC-type gene in the stem lineage of extant land plants (Henschel, et al. 2002; Kaufmann, et al. 2005; Gramzow and Theißen 2010). Consequently, MIKC-type genes from charophyte green algae, a grade of freshwater algae that represent land plants’ closest extant relatives, phylogenetically belong to neither of the two land-plant specific subfamilies (Nishiyama, et al. 2018). Instead charophyte MIKC-type genes likely constitute direct descendants of an ancestral MIKC-type gene, prior to the split into MIKC^C^- and MIKC*-type. We analysed the exon-intron structures of charophyte MIKC-type genes, for which genomic information is available, and indeed found the hypothetical exon-intron structure of an ancestral MIKC-type gene. The charophyte MIKC-type protein KnMADS1 from *Klebsormidium nitens* (Klebsormidiophyceae) is encoded by one MADS-domain exon, one I-domain exon, four K-domain exons and five C-terminal domain exons (Figure 3c). The first two K-domain exons of *KnMADS1* show high similarity in length and sequence to the first and second, as well as to the third and fourth K-domain exon of MIKC*-type genes, whereas the last K-domain exon of *KnMADS1* shows high similarity to the last two K-domain exons of MIKC^C^-type genes (Figure 3c). This observation corroborates the hypothesis that ancestral MIKC^C^- and MIKC*-type genes underwent exon duplications causing structural divergence of their K-domains.

To better comprehend which structural changes may have been brought about by the different exon duplications, we plotted the protein regions encoded by the different K-domain exons on the known crystal structure of the K-domain of the MIKC^C^-type protein SEP3 from *A. thaliana* (Puranik, et al. 2014). *SEP3* exon 3 (i.e. the first K-domain exon) encodes for the first (N-terminal) α-helix harbouring dimerisation interface 1. Exon 4 encodes for the kink region separating both K-domain helices and for the N-terminal half of the second α-helix harbouring dimerization interface 2. Exons 5 and 6 encode for the C-terminal half of the second α-helix comprising the tetramerisation interface (Figure 3d and e). Consequently, the duplication of the last K-domain exon of an ancestral MIKC^C^-type gene likely elongated the second (C-terminal) K-domain helix and thereby gave rise to the full-length tetramerisation interface found in extant MIKC^C^-type proteins. In contrast, the duplication of the first two K-domain exons of an ancestral MIKC*-type protein probably elongated the dimerization interface of the first (N-terminal) helix.

### Artificial exon deletion impedes FQC formation of MIKC^C^-type proteins

To investigate the functional importance of the last two K-domain exons for FQC formation of MIKC^C^-type proteins, we generated mutant versions of *PPM1*, *SmMADS3*, *CRM3*, and *SEP3*, lacking the sequence of either of the two hypothetically duplicated K-domain exons (exon 5 and exon 6, respectively), by mutagenesis PCR. The resulting mutant proteins PPM1ΔE5, PPM1ΔE6, SmMADS3ΔE5, SmMADS3ΔE6, CRM3ΔE5, CRM3ΔE6, SEP3ΔE5 and SEP3ΔE6 were expressed *in vitro* and tested for their ability to form FQCs in EMSA. In contrast to the wild type proteins, all exon deletion mutants, except for CRM3ΔE6, produced a fraction of intermediate electrophoretic mobility (DNA probe bound by two proteins) at low amounts of applied protein. With increasing protein amounts an additional fraction of low electrophoretic mobility (DNA probe bound by four proteins) occurred (Figure 4 and Supplementary Figure 3). By quantification of the signal intensities of the different fractions, FQC formation capabilities were estimated and expressed as *kd1/kd2*. Except for CRM3ΔE6, all mutated MIKC^C^-type proteins produced low cooperativity values, comparable to those determined for MIKC*-type proteins (Figure 4). This indicates no or very weak positive interaction between the two DNA-bound dimers and thus no ability for cooperative DNA-binding and FQC formation. CRM3ΔE6 showed an uncommon DNA-binding behaviour as it produced two fractions of low electrophoretic mobility. Furthermore, the bound and unbound fractions did not follow the usually observed sigmoidal increase and decrease, respectively (Figure 4f). However, as no signal of a DNA probe bound by only two proteins was observed, it appears likely that CRM3ΔE6 is still able to form FQCs.

**Figure 4:**
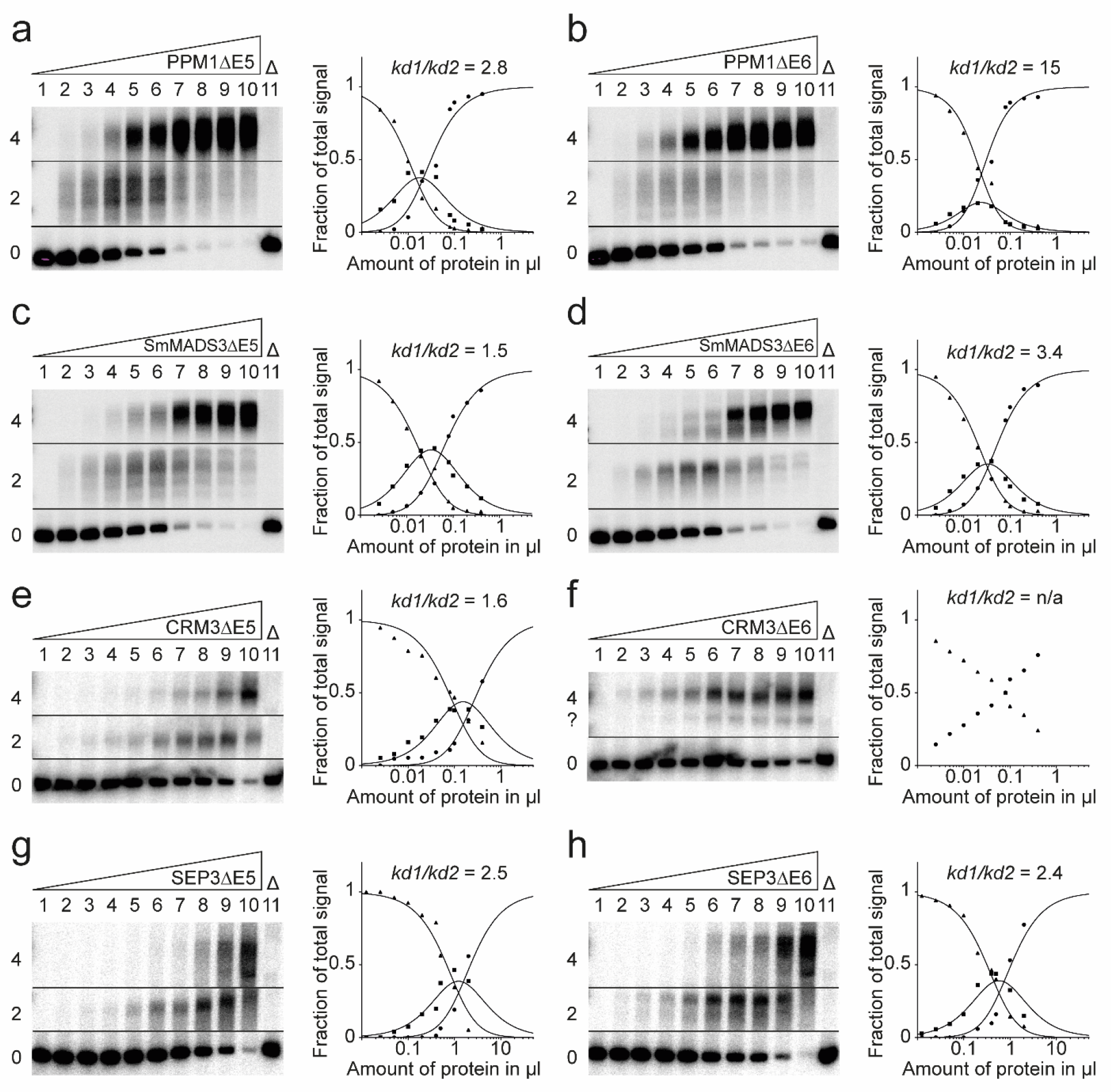
FQC formation capabilities of exon deletion mutants of the MIKC^C^-type proteins PPM1, SmMADS3, CRM3, and SEP3. Increasing amounts of *in vitro* transcribed/translated (a) PPM1ΔE5, (b) PPM1ΔE6, (c) SmMADS3ΔE5, (d) SmMADS3ΔE6, (e) CRM3ΔE5, (f) CRM3ΔE6, (g) SEP3ΔE5, and (h) SEP3ΔE6 were co-incubated together with constant amounts of radioactively labelled DNA probe carrying two SRF-type CArG-boxes. Up to three fractions of different electrophoretic mobility occur – a fraction of high electrophoretic mobility constituting unbound DNA probe (labelled with ‘0’), a fraction of intermediate electrophoretic mobility constituting a DNA probe bound by a single protein dimer (‘2’) and a fraction of low electrophoretic mobility constituting a DNA probe bound by four proteins (‘4’). Signal intensities of different fractions were measured and plotted against the amount of applied protein (triangles, free DNA; squares, DNA probe bound by two proteins; circles, DNA probe bound by four proteins). Graphs were fitted according to equations (1) to (3) described in Materials and Methods to eventually quantify and express FQC formation capabilities by *k_d1_/k_d2_*. Because (f) CRM3ΔE6 produced no signal of intermediate electrophoretic mobility constituting a DNA probe bound by a single protein dimer, *k_d1_/k_d2_* cannot be determined. As negative control 2 μl of *in vitro* transcription/translation mixture loaded with the empty pTNT plasmid were added to the binding reaction (‘Δ’). Applied amounts of *in vitro* transcription/translation products were (lane 1 to 10) 0, 0.0125, 0.025, 0.05, 0.1, 0.2, 0.4, 0.5, 1, and 2 μl, whereby PPM1ΔE5, PPM1ΔE6, SmMADS3ΔE5, SmMADS3ΔE6, CRM3ΔE5, and CRM3ΔE6 were prediluted 1:5 with BSA (10 mg ml^-1^).

### Artificial duplication of exon 6 enables KnMADS1 to form FQCs

To further study the functional implications of the exon duplications during early evolution of MIKC^C^- and MIKC*-type genes, we used the charophyte MIKC-type gene *KnMADS1* as a proxy for the ancestral state of land plant MIKC^C^- and MIKC*-type genes immediately after the split into both gene families. We generated three mutated versions of *KnMADS1* (Figure 5a). To mimic an early MIKC^C^-type gene, we duplicated exon 6 (*KnMADS1duplE6*) and in addition deleted exon 3 (*KnMADS1Δ3duplE6*), as exon 3 of *KnMADS1* is not found among extant MIKC^C^-type genes (Figure 2 and Figure 3c). To mimic an early MIKC*-type gene, we duplicated *KnMADS1* exons 3 and 4 (*KnMADS1duplE3E4*). KnMADS1 wild type and mutated proteins were produced *in vitro* and tested for their ability to form FQCs in EMSA. In accordance to our hypothesis that the duplication of the last K-domain exon facilitated FQC formation, KnMADS1 wild type protein produced very low cooperativity values and thus was unable to form FQCs under our experimental conditions (Figure 5b and Supplementary Figure 4a). In contrast, the two MIKC^C^-type mimicking mutants KnMADS1duplE6 and KnMADS1ΔE3duplE6 showed no signal of intermediate electrophoretic mobility (DNA probe bound by only two proteins), but instead bound to DNA immediately as tetramer, similar to the tested MIKC^C^-type proteins from Physcomitrium, Selaginella and Ceratopteris (Figure 5c,d and Supplementary Figure 4b,c). The MIKC*-type mimicking mutant KnMADS1duplE3E4 showed a DNA-binding behaviour similar to that of the KnMADS1 wild type protein, although the signal of intermediate electrophoretic mobility appeared weaker, resulting in slightly higher cooperativity values (Figure 5e and Supplementary Figure 4d).

**Figure 5:**
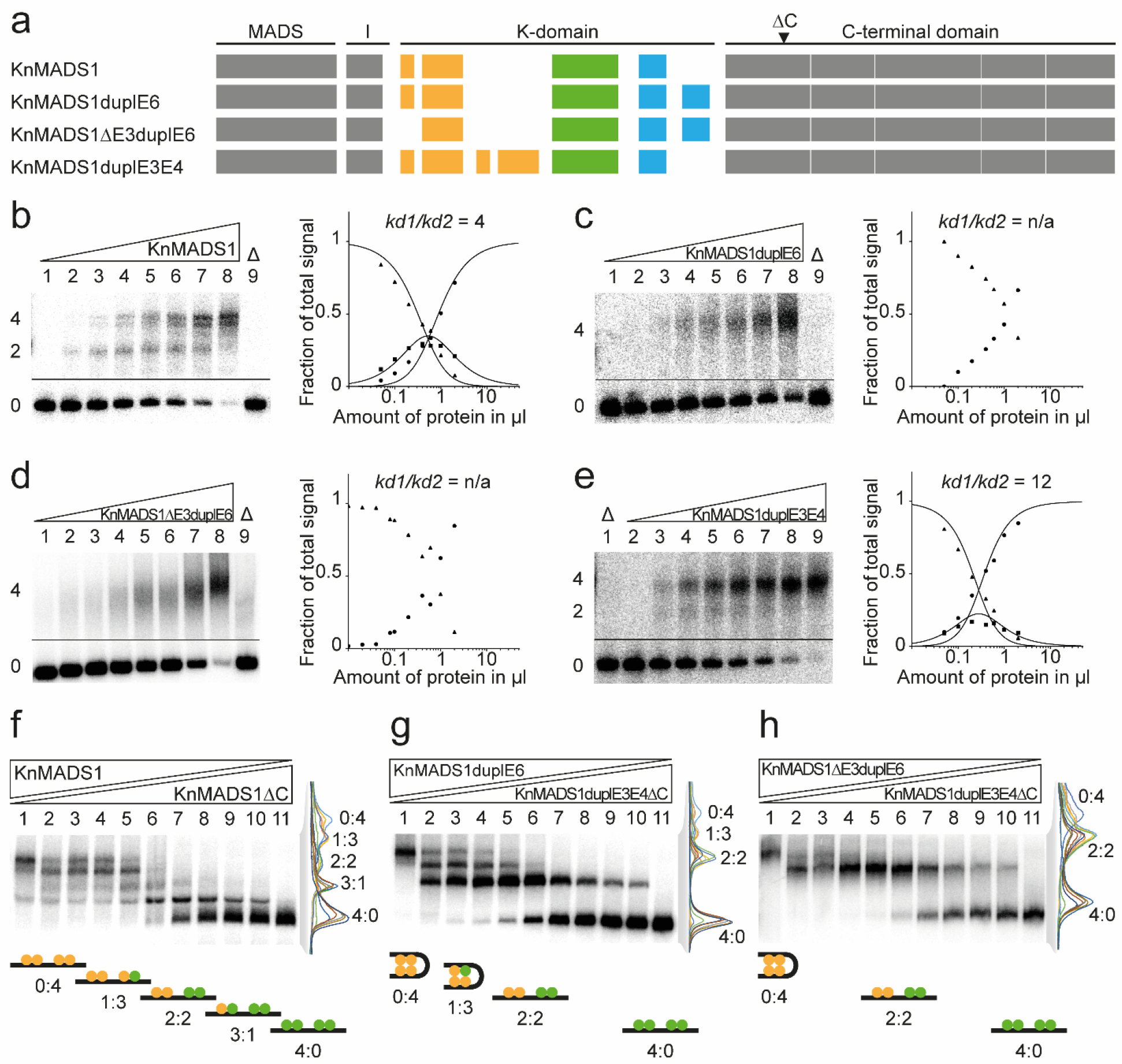
FQC formation capabilities of exon duplication and deletion mutants of the charophyte MIKC-type protein KnMADS1. (a) Exon-intron structure of KnMADS1 wild type and the mutated version KnMADS1duplE6, KnMADS1ΔE3duplE6, and KnMADS1duplE3E4. K-domain exons are colour-coded according to Figure 3c. Black triangle highlights the position at which the coding sequence was terminated to generate C-terminally truncated versions of KnMADS1 and KnMADS1duplE3E4. (b-e) Increasing amounts of *in vitro* transcribed/translated (b) KnMADS1, (c) KnMADS1duplE6, (d) KnMADS1ΔE3duplE6, and (e) KnMADS1duplE3E4 was co-incubated together with constant amounts of radioactively labelled DNA probe carrying two SRF-type CArG-boxes. Up to three fractions of different electrophoretic mobility occur – a fraction of high electrophoretic mobility constituting unbound DNA probe (labelled with ‘0’), a fraction of intermediate electrophoretic mobility constituting a DNA probe bound by a single protein dimer (‘2’) and a fraction of low electrophoretic mobility constituting a DNA probe bound by four proteins (‘4’). Signal intensities of different fractions were measured and plotted against the amount of applied protein (triangles, free DNA; squares, DNA probe bound by two proteins; circles, DNA probe bound by four proteins). Graphs were fitted according to equations (1) to (3) described in Materials and Methods to eventually quantify and express FQC formation capabilities by *k_d1_/k_d2_*. Because (c) KnMADS1duplE6 and (d) KnMADS1ΔE3duplE6 produced no signals of intermediate electrophoretic mobility constituting a DNA probe bound by a single protein dimer, *k_d1_/k_d2_* cannot be determined. (f) KnMADS1 wild type protein was co-expressed at different ratios with KnMADS1ΔC and co-incubated together with constant amounts of the same radioactively labelled DNA probe as in b-e. In total, five retarded fractions of different electrophoretic mobility occurred. An overly of measured signal intensities of the individual lanes is shown on the right. Each peak of the graph is labelled according to the ratio of wild type and truncated protein of the corresponding fraction. The cartoons below the gel illustrate the composition of the different fractions with full-length and truncated proteins shown in yellow and green, respectively. (g,h) To test for heteromeric interaction capabilities between KnMADS1duplE6, KnMADS1ΔE3duplE6, and KnMADS1duplE3E4, the same assay as in panel f was conducted using (g) KnMADS1duplE6 together with KnMADS1duplE3E4ΔC and (h) KnMADS1ΔE3duplE6 together with KnMADS1duplE3E4ΔC, respectively. As negative control 2 μl of *in vitro* transcription/translation mixture loaded with the empty pTNT plasmid were added to the binding reaction (‘Δ’). (b-e) Applied amounts of *in vitro* transcription/translation products were (lane 1 to 8) 0, 0.05, 0.1, 0.2, 0.4, 0.6, 1, and 2 μl. (f-h) 3 μl of *in vitro* transcription/translation product were applied to each lane. Ratios of both template plasmids used for *in vitro* transcription/translation were (lane 1 to 11): 0:1, 1:9, 1:7, 1:5, 1:3, 1:1, 3:1, 5:1, 7:1, 9:1, 1:0.

### Exon duplication mutants of KnMADS1 are unable to form heterodimers

Based on large-scale interaction-data of MIKC^C^- and MIKC*-type proteins from seed plants, both subfamilies form two mostly separated protein-protein interaction networks (de Folter, et al. 2005; Immink, et al. 2009; Smaczniak, Immink, Angenent, et al. 2012). We aimed to investigate, whether the structural divergence of the K-domains of MIKC^C^- and MIKC*-type proteins, that was brought about by the duplication of different K-domain exons, may impede interactions between members of both subfamilies. We generated C-terminal truncation mutants of *KnMADS1* wild type (*KnMADS1ΔC*) and *KnMADS1duplE3E4* (*KnMADS1duplE3E4ΔC*) in order to be able to differentiate heteromeric complexes of different composition in EMSA due to their different size. When we co-incubated variable amounts of KnMADS1 and KnMADS1ΔC together with labelled DNA probe, five retarded fractions of different electrophoretic mobility occurred, demonstrating that each DNA-bound KnMADS1 wild type protein can be substituted by a C-terminally truncated version (Figure 5f and Supplementary Figure 4e). When we conducted the same experiment using variable amounts of KnMADS1duplE6 and KnMADS1duplE3E4ΔC, only four retarded fractions of different electrophoretic mobility occurred (Figure 5g and Supplementary Figure 4f). The fraction representing a DNA probe bound by three KnMADS1duplE3E4ΔC proteins and one KnMADS1duplE6 protein (fraction labelled with ‘3:1’ in Figure 5f) was completely absent. When we co-incubated variable amounts of KnMADS1ΔE3duplE6 and KnMADS1duplE3E4ΔC, only three retarded fractions occurred, likely representing a DNA probe bound by four KnMADS1ΔE3duplE6 proteins (labelled with ‘0:4’), by two KnMADS1ΔE3duplE6 proteins and two KnMADS1duplE3E4ΔC proteins (‘2:2’) and by four KnMADS1duplE3E4ΔC proteins (‘4:0’), respectively (Figure 5h and Supplementary Figure 4g). No DNA probe bound by three KnMADS1duplE3E4ΔC proteins and one KnMADS1ΔE3duplE6 protein or by one KnMADS1duplE3E4ΔC protein and three KnMADS1ΔE3duplE6 proteins was observed. Consequently, the duplication of exon 3 and 4 of one partner and the duplication of exon 6 (and deletion of exon 3) of the other partner seems to impede heterodimer formation of KnMADS1. Therefore, it appears plausible that the structural divergence of K-domains of MIKC^C^- and MIKC*- type proteins, that was caused by the different exon duplications, prevents heteromeric interactions between members of both subfamilies.

### Some charophyte MIKC-type proteins may independently have gained the ability to form FQCs

Besides KnMADS1, we cloned and tested a number of other charophyte MIKC-type proteins for their ability to form FQCs. Similar to KnMADS1, also CaMADS1 from *Chlorokybus atmophyticus* (Chlorokybophyceae) and CglMADS1 from *Chaetosphaeridium globosum* (Coleochaetophyceae) produced clear signals of intermediate electrophoretic mobility (i.e. binding of a single protein dimer) resulting in low cooperativity values indicating no FQC formation (Figure 6a,b and Supplementary Figure 5a,b). In contrast, ZspMADS1 from *Zygnema sp*. (Zygnematophyceae) produced only very weak signals of intermediate electrophoretic mobility resulting in high cooperativity values denoting strong FQC formation capabilities (Figure 6c and Supplementary Figure 5c). All three tested proteins from the genus Coleochaete (Coleochaetophyceae), CiMADS1 from *Coleochaete irregularis*, CoMADS1 from *Coleochaete orbicularis*, and CsMADS1 from *Coleochaete scutata*, showed no signals of intermediate electrophoretic mobility at all, indicating no binding of single dimers and thus very strong FQC formation (Figure 6d-f, Supplementary Figure 5d-f). All experimentally investigated charophyte MIKC-type genes show no duplication of the last K-domain exon. However, the last K-domain exon of charophyte MIKC-type genes already encodes for the N-terminal half of the tetramerization interface found in land plant MIKC^C^-type proteins. Therefore, it appears plausible, that to a certain extend already a shorter C-terminal K-domain helix is sufficient to mediate MTF tetramerisation.

**Figure 6:**
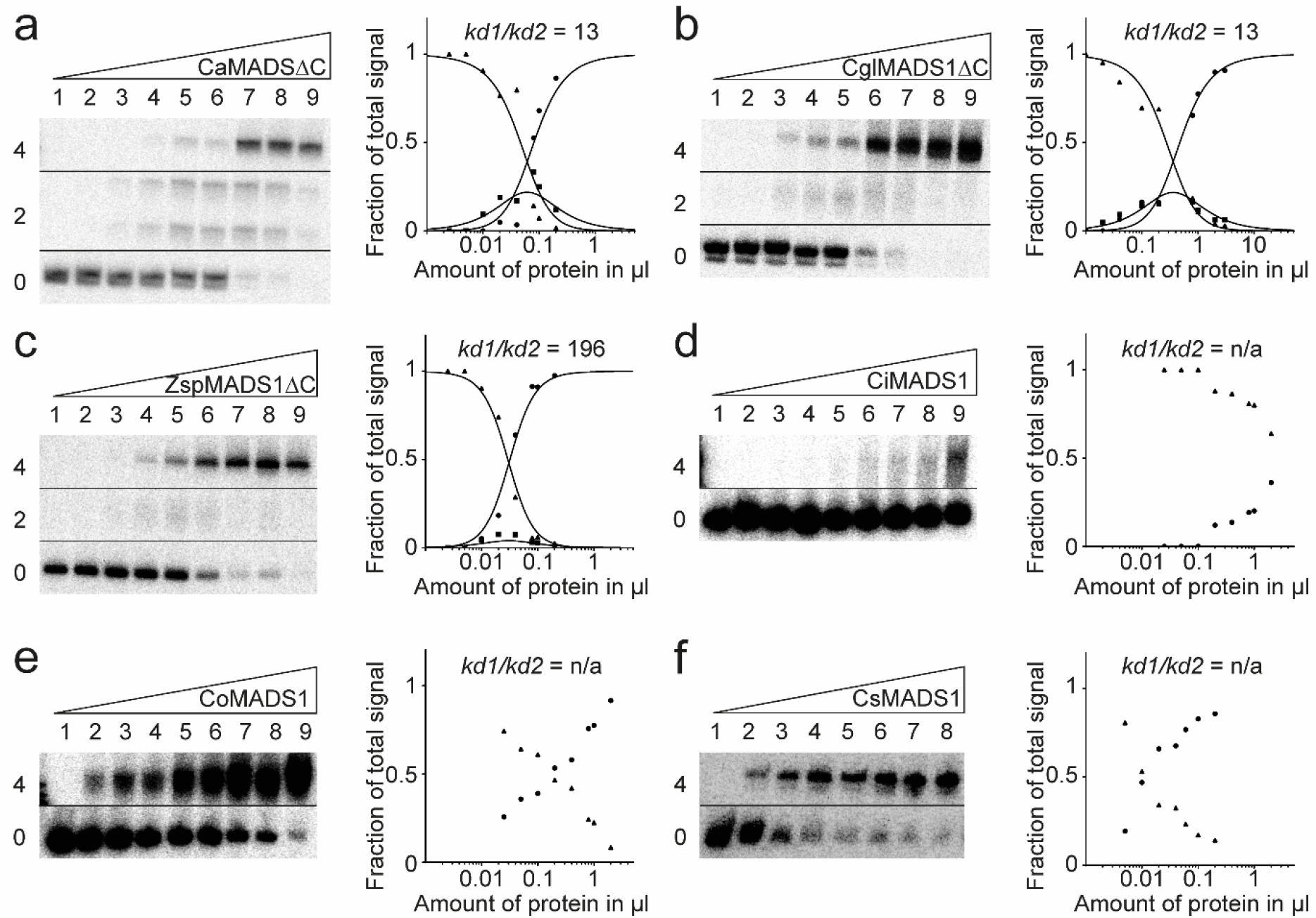
FQC formation capabilities of the charophyte MIKC-type proteins CaMADS1, CglMADS1, ZspMADS1, CiMADS1, CoMADS1, and CsMADS1. Increasing amounts of *in vitro* transcribed/translated (a) CaMADS1ΔC, (b) CglMADS1ΔC, (c) ZspMADS1ΔC, (d) CiMADS1, (e) CoMADS1, and (f) CsMADS1, were co-incubated together with constant amounts of radioactively labelled DNA probe carrying two N10-type CArG-boxes. Up to three fractions of different electrophoretic mobility occur – a fraction of high electrophoretic mobility constituting unbound DNA probe (labelled with ‘0’), a fraction of intermediate electrophoretic mobility constituting a DNA probe bound by a single protein dimer (‘2’) and a fraction of low electrophoretic mobility constituting a DNA probe bound by four proteins (‘4’). Signal intensities of different fractions were measured and plotted against the amount of applied protein (triangles, free DNA; squares, DNA probe bound by two proteins; circles, DNA probe bound by four proteins). Graphs were fitted according to equations (1) to (3) described in Materials and Methods to eventually quantify and express FQC formation capabilities by *k_d1_/k_d2_*. Because (d) CiMADS1, (e) CoMADS1, and (f) CsMADS1 produced no signal of intermediate electrophoretic mobility constituting a DNA probe bound by a single protein dimer, *k_d1_/k_d2_* cannot be determined. In case of (a) CaMADS1 a double band was observed for the fraction of intermediate electrophoretic mobility, likely caused by different conformations of the DNA-bound protein dimer. Because CaMADS1, CglMADS1, and ZspMADS1 full-length proteins comprise very long C-terminal domains, resulting in low DNA-binding affinities and blurry band, C-terminally truncated mutants were used instead. As negative control 2 μl of *in vitro* transcription/translation mixture loaded with the empty pTNT plasmid were added to the binding reaction (‘Δ’). Applied amounts of *in vitro* transcription/translation products were (a, c-e) (lane 1 to 9) 0, 0.025, 0.05, 0.1, 0.2, 0.4, 0.8, 1, and 2 μl, whereby CaMADS1ΔC and ZspMADS1ΔC were prediluted 1:10 with BSA (10 mg ml^-1^), (b) (lane 1 to 9) 0.02, 0.04, 0.1, 0.2, 0.4, 0.8, 1, 2, and 3 μl, (f) (lane 1 to 8) 0, 0.005, 0.01, 0.02, 0.04, 0.06, 0.1, and 0.2 μl.

## DISCUSSION

Floral-quartet-like complexes (FQCs) represent a unique system of gene regulation involving heterotetramers of MIKC^C^-type MADS-domain transcription factors (MTFs). FQCs control important developmental processes in plants, but how they originated remained unknown for decades. Here we present data clarifying the evolutionary origin of FQCs.

Like many other proteins, MTFs bind with sufficient strength to specific DNA sequences only as dimers, a feature that they share also with all other MADS-domain proteins (Shore and Sharrocks 1995; Amoutzias, et al. 2008). In addition, however, many MTFs constitute also tetrameric complexes. These tetrameric MTFs recognise their target genes by binding to two CArG-boxes, involving DNA-loop formation between the binding sites (Egea-Cortines, et al. 1999; Theißen 2001; Theißen and Saedler 2001; Theißen, et al. 2016). Tetramerisation of regulatory proteins, and DNA-binding on two cis-regulatory elements involving DNA looping, is well known from bacterial repressors and activators, such as the lac repressor and lambda repressor/activator (Hochschild and Ptashne 1986; Oehler, et al. 1990; Lewis 2005). In these cases, protein-protein interactions between protein dimers provide Gibbs free energy (ΔG°) in addition to that available by protein-DNA interactions alone, and hence lead to a cooperative formation of tetramers bound to DNA. Consequently, a switch-like on-off interaction of the regulatory proteins with their target genes occurs (Hochschild and Ptashne 1986).

Cooperative binding has also been demonstrated for MTFs in FQCs (Melzer, et al. 2009; Jetha, et al. 2014; Rümpler, et al. 2018). The proximate (mechanistic) and ultimate (evolutionary) mechanisms that may have led to the origin and maintenance of MTF tetramerisation and FQC formation have already been discussed elsewhere (Theißen, et al. 2016). Briefly, like in case of viral and bacterial activators and repressors, cooperative formation of MTFs could lead to a sharp transcriptional response of target genes. Even small increases of MTF concentrations may lead to drastic changes in the regulatory response of target genes, allowing for a switch-like, all-or-nothing kind of regulation of target genes. Since MTFs act as genetic switches that control discrete developmental processes - the establishment of different kinds of tissue or organ identities is a good case in point -, cooperativity could well be one crucial mechanism that transforms the quantitative nature of molecular interactions between MTFs and DNA into discrete phenotypic outputs (Theißen and Melzer 2007; Kaufmann, et al. 2010). In case of e.g. organ identity genes the biological relevance of this behaviour is quite plausible, since it leads to the development of organs with unambiguous identities (such as stamens and carpels) and avoids the formation of chimeric organs. Accordingly, it has been hypothesized that the ability of MIKC^C^-type proteins to establish and diversify sharp developmental switches may have facilitated the evolution of different tissues and organs as required during evolution of the increasingly complex body plans of plants on land (Theißen, et al. 2016; Nishiyama, et al. 2018).

We consider tetramerisation of MTFs and FQC formation an important evolutionary novelty in gene regulation. It has long been known that MADS-domain proteins act in multimeric (often called „ternary“) complexes. In cases other than MIKC^C^-type proteins, however, dimers of MADS-domain proteins interact with non-MADS-domain proteins. Well-characterized examples are the different complexes of the general yeast transcription factor MINICHROMOSOME MAINTENANCE 1 (MCM1), some of which involve e.g. the homeodomain protein α2 and the HMG-domain protein α2, and the animal SERUM RESPONSE FACTOR (SRF) that interacts among others with members of the myocardin family of transcription factors and with ternary complex factors (Shore and Sharrocks 1995; Messenguy and Dubois 2003; Tsong, et al. 2006; Onuh and Qiu 2021). Tetrameric complexes composed exclusively of MADS-domain proteins encoded by the same or paralogous genes appear to be unique to MTFs in plants, however.

Heterotetramerisation of paralogous MTFs represents not only a regulatory system with important functions in flowering plants, but, by inference, very likely also in all other land plants (embryophytes). Hence tetramerisation of MTFs and cooperative formation of FQCs is of considerable biological and even agronomic interest. Surprisingly little has been known about its evolutionary origin, however. It has been shown that in addition to MIKC^C^-type proteins of flowering plants also some MIKC^C^-type proteins of extant gymnosperms are able to form FQCs (Wang, et al. 2010; Ruelens, et al. 2017). Since the lineages that led to extant gymnosperms and angiosperms separated roughly about 300 million years ago, this pushes the origin of FQC formation accordingly. Corresponding information about non-seed plants, and about MIKC*-type proteins from any plants, was lacking so far. It remains unclear, therefore, when (in time) and where (in which lineage) tetramerisation of MTFs actually originated. Here we demonstrate that FQC formation has much deeper roots than seed plants, down to streptophyte green algae, which are close relatives of land plants. FQC formation had therefore been established at least 450 – 500 million years ago already.

Almost twenty years ago two hypothetical scenarios have been proposed to explain the causal link between the K-domain and MTF tetramerisation (Kaufmann, et al. 2005). One model has it that the K-domain was initially just involved in protein dimerisation and is hence more ancient than tetramerisation. Only later tetramerisation may have been acquired as a second function of the K-domain. Alternatively, the K-domain might have had a primary role in tetramerisation right from the beginning, and no or only a minor role in dimerization. Under this assumption, the MTFs and tetramerisation would have occurred simultaneously.

Based on the data presented here, we now can draw a clear picture of the molecular changes that facilitated MTF tetramerisation (Figure 7). In the stem group of extant streptophytes, a K-domain encoding sequence originated downstream of the MADS-box, by a yet unknown mechanism. In its ancestral state, the K-domain was likely encoded by 4 exons and probably already constituted a fold similar to that determined for the MIKC^C^-type protein SEP3 (Puranik, et al. 2014), comprising helix 1, a rigid kink and helix 2. This ancestral type of K-domain (as still retained in some charophyte green algae) may had already been able to facilitate tetramerisation to a certain extent, but doesn’t do so in all cases (Figure 6), so that it may have primarily originated as an additional dimerization domain. In the stem group of extant land plants, a gene duplication of an ancestral MIKC-type gene occurred (Figure 7b). In the lineage of one of the copies, the first two K-domain exons got duplicated, generating MIKC*-type genes as has already been hypothesised before (Figure 3b,c) (Kwantes, et al. 2012). In the lineage of the second copy, the first K-domain exon likely got lost, whereas the last K-domain exon was duplicated, generating MIKC^C^-type genes (Figure 3a,c). By the duplication of the last K-domain exon, helix 2 of the K-domain got elongated, now constituting a strong tetramerisation interface (Figure 3d,e). Since no MIKC-type genes encoding ancestral K-domains have been found in land plant genomes yet, it appears very likely that such genes have been lost in the stem group of extant land plants, even though other scenarios currently cannot be completely ruled out. Ancestral MTFs may have negatively interfered with the assembly or function of transcription factor complexes once more ‘advanced’ MIKC^C^-type and MIKC*-type proteins had originated.

**Figure 7:**
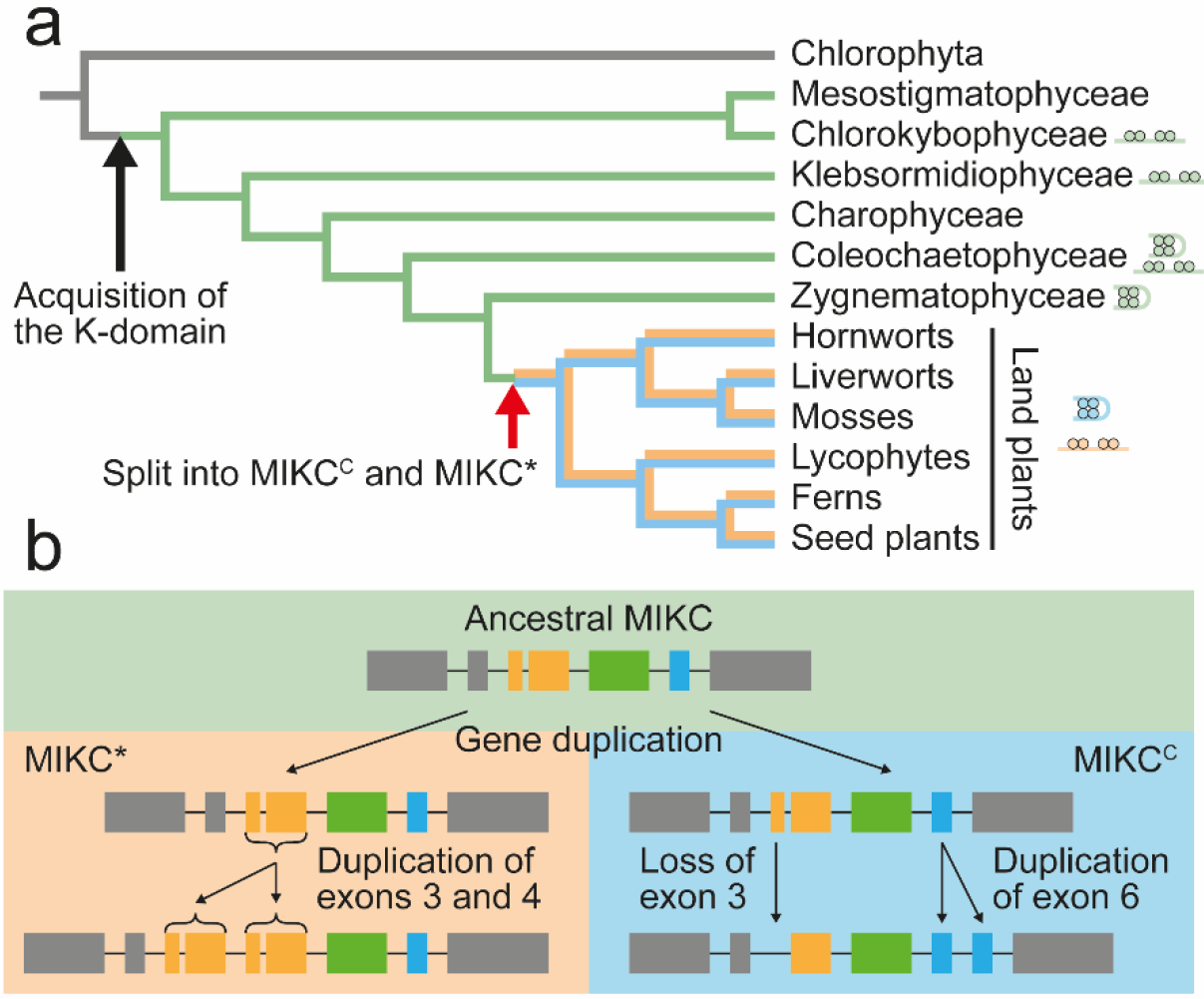
Hypothesized mode of MTF evolution. (a) Simplified phylogenetic tree of green plants with highlighted major evolutionary changes of Type II MADS-box genes (branching according to Wickett, et al. (2014)). In the stem group of extant streptophytes (charophytes + land plants), an ancestral Type II MADS-box gene (illustrated by grey branch color) acquired the K-domain, giving rise to the eponymous ‘MIKC’ domain architecture of the encoded MTFs. In the stem group of extant land plants, a gene duplication of an ancestral MIKC-type gene (illustrated by green branch color) gave rise to the two, land plant specific, MTF subfamilies MIKC^C^ (blue branch color) and MIKC* (orange branch color). Red arrow depicts the time point, when the evolutionary changes shown in panel b occurred. Cartoons of FQCs and DNA-bound MTF dimers indicate presence or absence, respectively, of FQC formation capabilities of MTFs in the respective plant lineages. (b) Following the gene duplication of an ancestral MIKC-type gene (green background), the first two K-domain exons got duplicated in the gene lineage of MIKC*-type genes (orange background), whereas the first K-domain exon got lost and the last K-domain exon got duplicated in the gene lineage of MIKC^C^-type genes (blue background).**Figure 3**

During the diversification of land plants into bryophytes (hornworts, liverworts, mosses), lycophytes, monilophytes (ferns and their allies) and spermatophytes (gymnosperms and angiosperms), gene duplications and sequence diversifications led to a moderate increase in the number of MIKC*-type genes and a strong increase in the number of MIKC^C^-type genes (Gramzow and Theißen 2010; Thangavel and Nayar 2018).

In seed plants, MIKC^C^-type genes are almost exclusively involved in the control of sporophyte developmental processes (Smaczniak, Immink, Angenent, et al. 2012). Given that the sporophyte became the dominant life phase during land plant evolution, it is tempting to speculate that the gain of tetramerisation in the MIKC^C^-type lineage, the strong increase in the number of MIKC^C^-type genes during land plant evolution, and the gain in importance of the sporophyte life phase were causally linked. MIKC^C^-type genes might have been co-opted to control developmental processes of the increasingly complex sporophyte, because they were able to cooperatively constitute FQCs in different combinations and therefore bore a higher combinatorial potential than MIKC*-type genes. The ability of MIKC^C^-type proteins to constitute FQCs may have enabled the control of numerous different tissues and organs and hence may have facilitated the origin of land plant diversity from the species to the body plan level.

Large scale interaction data of MIKC^C^- and MIKC*-type proteins from flowering plants have shown, that both protein subfamilies form two largely independent interaction networks (de Folter, et al. 2005; Immink, et al. 2009; Gramzow and Theißen 2010). Based on our data, it appears likely that the structural changes within the K-domain caused by duplications of different exons, laid the foundation for this functional separation and the subsequent evolution of independent gene regulatory networks.

## MATERIAL AND METHODS

### Cloning procedures and site-directed mutagenesis

The plasmid for *in vitro* transcription/translation of *A. thaliana SEP3* has been generated previously (Melzer, et al. 2009). Coding sequences of *Zygnema sp. ZspMADS1* (1kP: STKJ-2019062), *C. globosum CglMADS1* (1kP: DRGY-2036441), Co. scutata *CsMADS1* (GeneBank: AB035568.1), Co. irregularis *CiMADS1* (1kP:QPDY-2028842), Co. orbicularis *CoMADS1* (GBSL01006298.1), *K. nitens KnMADS1* (GeneBank: DF237509; locus tag KFL_005600030), *Ch. atmophyticus CaMADS1* (1kP:AZZW-2004234), *P. patens PPM1* (GeneBank: AF150931_1) and *PPM4* (GeneBank: AY509529.1), *S. moellendorffii SmMADS2* (NCBI Ref: XM_002974738.1) and *SmMADS3* (NCBI Ref: XM_002984875.1), *C. richardii CRM3* (GeneBank: Y08239_1), *CRM13* (GeneBank: FM995267.1), *CRM14* (GeneBank: FM995269.1), *CRM15* (GeneBank: FM995271.1), and *CRM16* (GeneBank: FM995273.1), as well as *A. thaliana AGL66* (NCBI Ref: NM_106447.4) and *AGL104* (NCBI Ref: NM_102063.3) were codon optimized for *Oryctolagus cuniculus*, synthesized via the GeneArt gene synthesis service (Thermo Fisher Scientific) and cloned into pTNT (Promega) using *Eco*RI and *Sal*I restriction sites. Plasmids for *in vitro* transcription/translation of C-terminally truncated proteins, exon duplication and exon deletion mutants were generated by site-directed mutagenesis PCR following the manufacturer’s instructions of the Q5 Site-Directed Mutagenesis Kit (New England Biolabs). To generate the plasmids for *in vitro* transcription/translation of the fusion proteins SmMADS3-GFP and CRM3-GFP, a *Hind*III restriction site was introduced into plasmids pTNT-SmMADS3 and pTNT-CRM3 by site-directed mutagenesis substituting the stop codons of SmMADS3 and CRM3. Subsequently, the coding sequence of the enhanced GFP gene mGFP6 was PCR amplified from pGreenII-35S::mGFP6 (Hellens, et al. 2000) while adding *Hind*III restriction sites at the 3’ and 5’ end. The purified PCR product was cloned into pTNT-SmMADS3 and pTNT-CRM3 using the introduced *Hind*III restriction site.

### Generation of DNA probes

The DNA probes for electrophoretic mobility shift assay (EMSA) have been generated essentially as described by Melzer, et al. (2009). Two different DNA probes were used in this study (Supplementary Table 1). For FQC-formation assays of MIKC^C^-type proteins, a 151-bp DNA probe was used that carried two SRF-type CArG-boxes of the sequence 5’-CCAAATAAGG-3’ in a distance of 63 bp. For FQC-formation assays of MIKC*-type proteins, the same DNA probe was used except that the SRF-type CArG-boxes were substituted by N10-type CArG-boxes of the sequence 5’-CTATATATAG-3’. The DNA probes were radioactively labelled with [α-P^32^] dATP by a Klenow fill-in reaction of 5’-overhangs.

### *In vitro* transcription/translation and electrophoretic mobility shift assay

Proteins were produced *in vitro* via the TNT SP6 Quick Coupled Transcription/Translation System (Promega). If two proteins were to be tested together in a single binding reaction, the proteins were co-expressed by loading the transcription/translation system with template plasmids of both proteins at the desired ratio. EMSA was performed essentially as described by Melzer, et al. (2009). The composition of the protein-DNA binding reaction buffer was essentially as described by Egea-Cortines, et al. (1999) with final concentrations of 1.6 mM EDTA, 10.3 mM HEPES, 1mM DTT, 1.3 mM Spermidine hydrochloride, 33.3 ng μl^-1^, Poly dI/dC, 2.5 % CHAPS, 4,3 % glycerol, and 5.2 to 7.7 ng μl^-1^ BSA. To test for FQC formation capabilities, 0.1 ng of radioactively labelled DNA probe was co-incubated with variable amounts of *in vitro* transcribed/translated protein ranging from 0.0025 to 2 μl. To test for complex stoichiometry and heteromeric interactions, 0.1 ng of radioactively labelled DNA probe was co-incubated with 3 μl of *in vitro* transcription/translation reaction product that had been loaded with both template plasmids at the indicated ratios: 0:1, 1:9, 1:7, 1:5, 1:3, 1:1, 3:1, 5:1, 7:1, 9:1, 1:0.

### Rationale of our FQC formation assay and quantification of FQC-formation capabilities

In this study, we analyse FQC formation of MTFs via a previously EMSA. As has been explained in detail elsewhere, our assay allows us to discriminate between two MTF dimers that individually bind to two adjacent CArG-boxes, and a single MTF tetramer that simultaneously binds to both CArG-boxes while looping the DNA in between and thus forming an FQC (Melzer and Theißen 2009; Melzer, et al. 2009). FQC formation is indicated by a high cooperativity of four MTFs binding to specific DNA sequences.

In brief, we incubate increasing amounts of the MTF of interest together with a constant amount of a radioactively labelled DNA probe carrying two identical CArG-boxes. In case of an MTF incapable of forming FQCs, dimers of MTFs will independently bind with the same affinity to both CArG-boxes. In other words, the affinity of the first dimer, binding to an unbound DNA probe, is equal to the affinity of the second dimer, binding to a DNA probe at which one binding site is already occupied. In contrast, in case of an MTF capable of forming FQCs, the two DNA-binding dimers positively interact with each other by forming a tetramer. Due to this positive interaction between both dimers, the affinity of the second dimer, that binds to a DNA probe at which one binding site is already occupied, is increased, compared to the DNA-binding affinity of the first dimer. This increased DNA-binding affinity of the second dimer, shifts the equilibrium of the binding reaction towards a DNA probe with two occupied binding sites. As a consequence, MTFs that are capable of forming FQCs produce weaker or no signals of a DNA probe bound by only one dimer than MTFs incapable of forming FQCs.

The signal intensities of the three fractions of different electrophoretic mobility (unbound DNA probe, DNA probe bound by two proteins, and DNA probe bound by four proteins) were quantified using Multi Gauge 3.1 (Fujifilm). Quantification of FQC-formation capabilities was performed by non-linear regression to equations described by Melzer, et al. (2009) and Senear and Brenowitz (1991). In brief, if the relative concentration of unbound DNA probe [Y0], DNA probe bound by two proteins [Y2], and DNA probe bound by four proteins [Y4] are described as a function of the amount of applied protein [P2]:

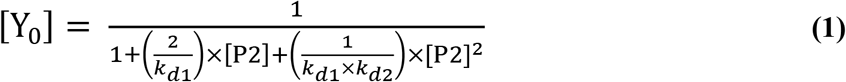

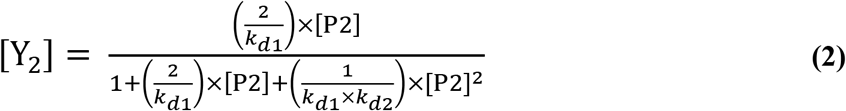

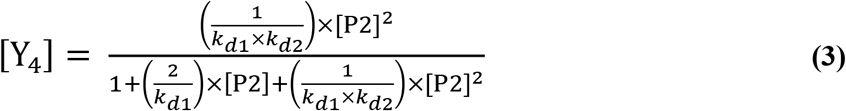

*k_d1_* and *k_d2_* are the dissociation constants for binding of a protein dimer to an unbound DNA probe and binding of a second protein dimer to a DNA probe where one binding site is already occupied, respectively. *kd1* and *kd2* were estimated by non-linear regression to equations (1) to (3) using GraphPad Prism 9 (GraphPad Software). The ability to form FQCs can subsequently be expressed by dividing *kd1* by *kd2*.

### Exon homology analysis and ancestral character state reconstruction

Coding sequences of all MIKC^C^- and MIKC*-type genes from *A. thaliana, C. richardii, S. moellendorffii* and *P. patens* were translated into protein sequences, aligned with Mafft applying default settings (Katoh and Standley 2013), and subsequently back-translated into a codon alignment using RevTrans (Wernersson and Pedersen 2003). Exon borders were determined by aligning coding sequences to the corresponding genomic sequences using Splign (Kapustin, et al. 2008). Information about sequence similarity and exon borders were combined into a graphical presentation utilizing a customized perl script previously described by Hoffmeier, et al. (2018). This way homologous exons were identified and manually aligned. Exons were colour-coded, depending on the protein domain they encode for. Exons were defined as K-domain exons if they were homologous to exons encoding for the K-domain of SEP3 from *A. thaliana*, for which the crystal structure has been determined (Puranik, et al. 2014). The codon alignments (including accession numbers for all genes) used for generation of Figure 2 and Figure 3 are given in Supplementary Data 1 and Supplementary Data 2, respectively.

Ancestral character states were reconstructed as described by Rümpler, et al. (2015) using Mesquite 3.7 (http://www.mesquiteproject.org).

## Supplementary Materials

**Supplementary Table 1:** Sequences of DNA probes used in EMSA.

**Supplementary Figure 1.** Original gel images of Figure 1a-f.

**Supplementary Figure 2:** Original gel images of Figure 1g-l.

**Supplementary Figure 3:** Original gel images of Figure 4.

**Supplementary Figure 4:** Original gel images of Figure 5.

**Supplementary Figure 5:** Original gel images of Figure 6.

**Supplementary Data 1:** Alignment used for the exon homology analysis of Figure 2.

**Supplementary Data 2:** Alignment used for the exon duplication analysis of Figure 3.

## Acknowledgements

We thank Felix Althoff and Sabine Zachgo (Osnabrück) for fruitful discussions about MADS-box genes on MAdLand, and to Stefan Rensing (Marburg/Freiburg) and Jan de Vries (Göttingen) for helpful information and stimulating discussions about diverse Charophyte genomes. Many thanks also to Philipp Gehlhaar for some experiments during initial stages of the project.

## Authors’ contributions

GT and FR initiated, conceived and supervised the project; FR, CT, CG, and MB generated the constructs for *in vitro* transcription/translation and conducted the EMSA experiments; CT and LG compiled the sequence collection and LG conducted the exon homology analysis; FR prepared figures; FR and GT wrote the manuscript; all authors read and approved the submitted version of the manuscript.

