## Supplementary Materials for "The origin of floral quartet formation - Ancient exon duplications shaped the evolution of MIKC-type MADS-domain transcription factor interactions"

**Supplementary Table 1:** Sequences of DNA probes used in EMSA (shown 5’ to 3’). CArG-boxes are highlighted in bold.

>SRF-type_CArG_box_probe

CTCGAGGTCGGAAATTTAATTATATT**CCAAATAAGG**AAAGTATGGAACGTTCGACGGTATCGATA
AGCTTGATGAAATTTAATTATATT**CCAAATAAGG**AAAGTATGGAACGTTATCGAATTCCTGCAGC
CCGGGGGATCCACTAGTTCTAGA

>N10-type_CArG_box_probe

CTCGAGGTCGGAAATTTAATTATATT**CTATATATAG**AAAGTATGGAACGTTCGACGGTATCGATA
AGCTTGATGAAATTTAATTATATT**CTATATATAG**AAAGTATGGAACGTTATCGAATTCCTGCAGC
CCGGGGGATCCACTAGTTCTAGA

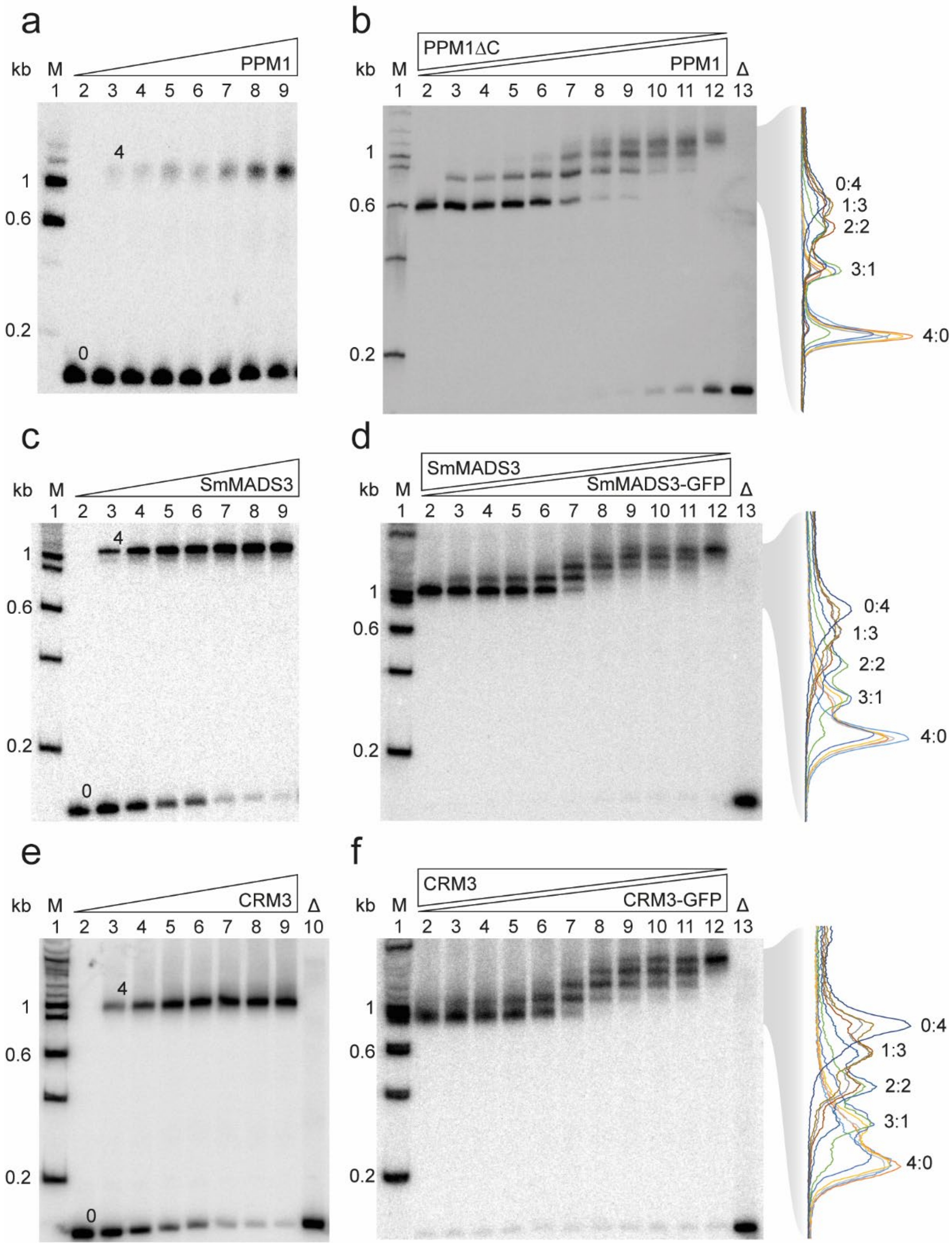


**Supplementary Figure 1. Original gel images of Figure 1a-f.** See figure legend of Figure 1. For size comparison radioactively labelled 100 bp DNA ladder (NEB) was added (‘M’).

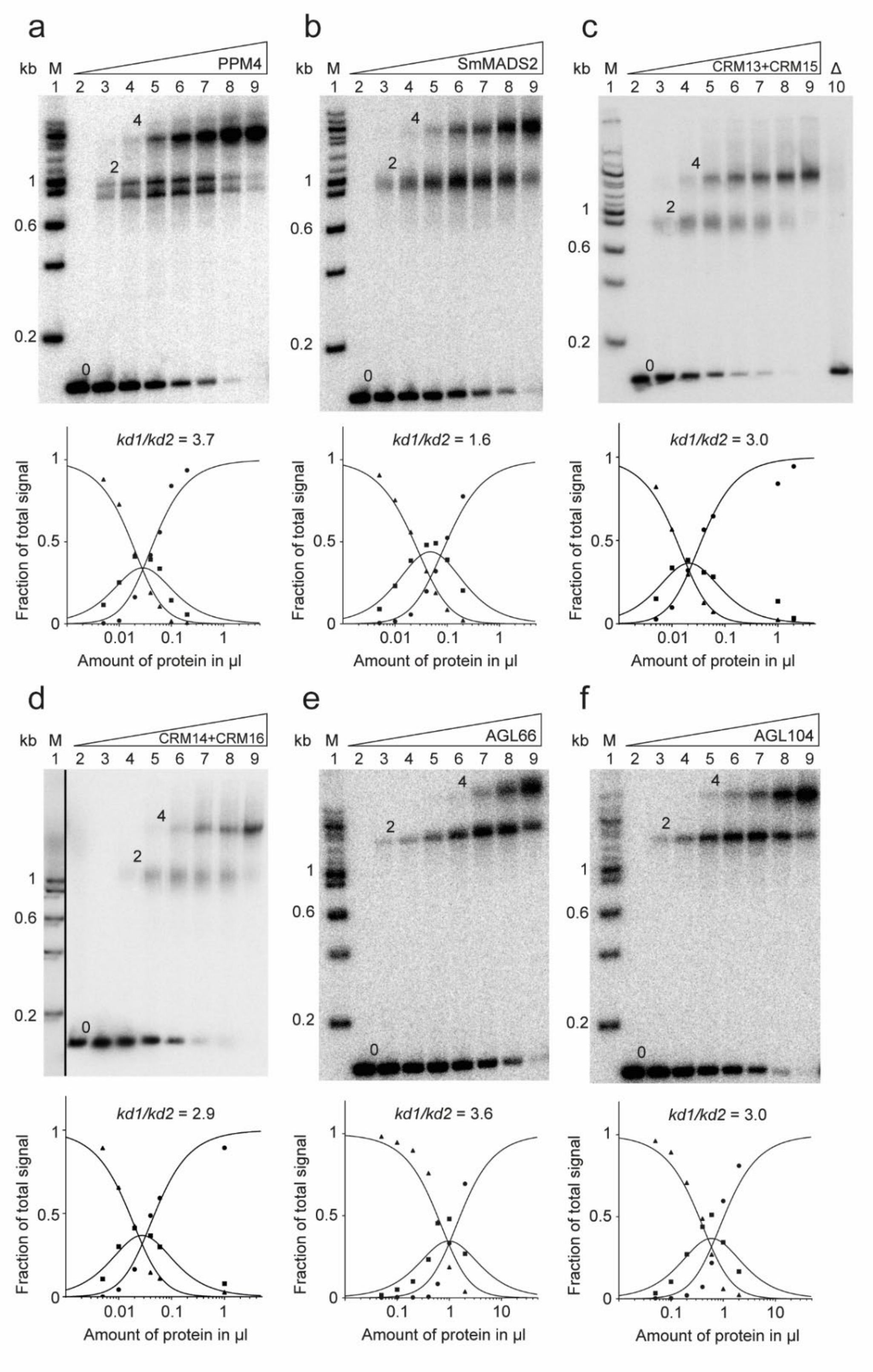


**Supplementary Figure 2: Original gel images of Figure 1g-l.** See figure legend of Figure 1. For size comparison radioactively labelled 100 bp DNA ladder (NEB) was added ('M').

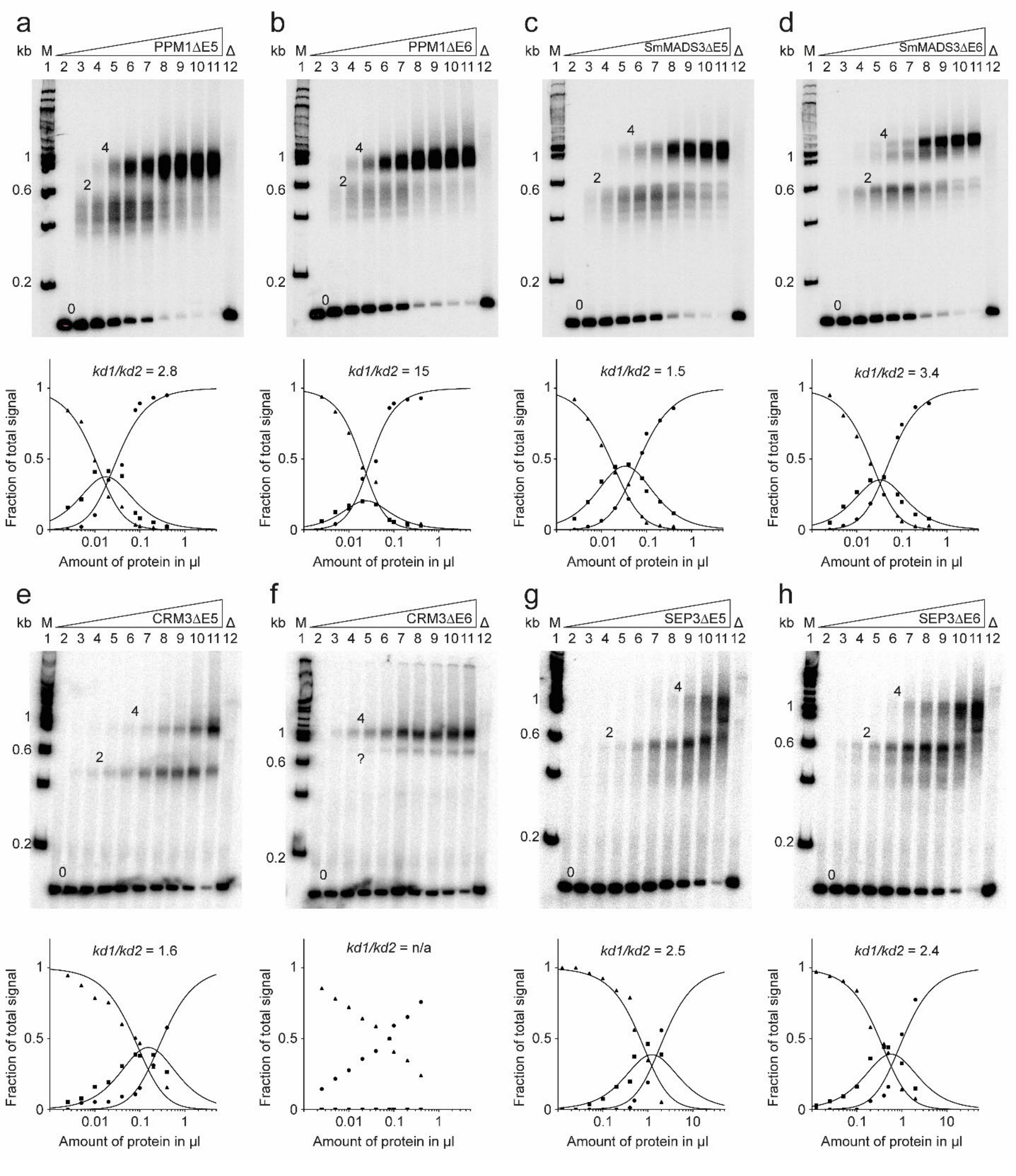


**Supplementary Figure 3: Original gel images of Figure 4.** See figure legend of Figure 4. For size comparison radioactively labelled 100 bp DNA ladder (NEB) was added ('M').

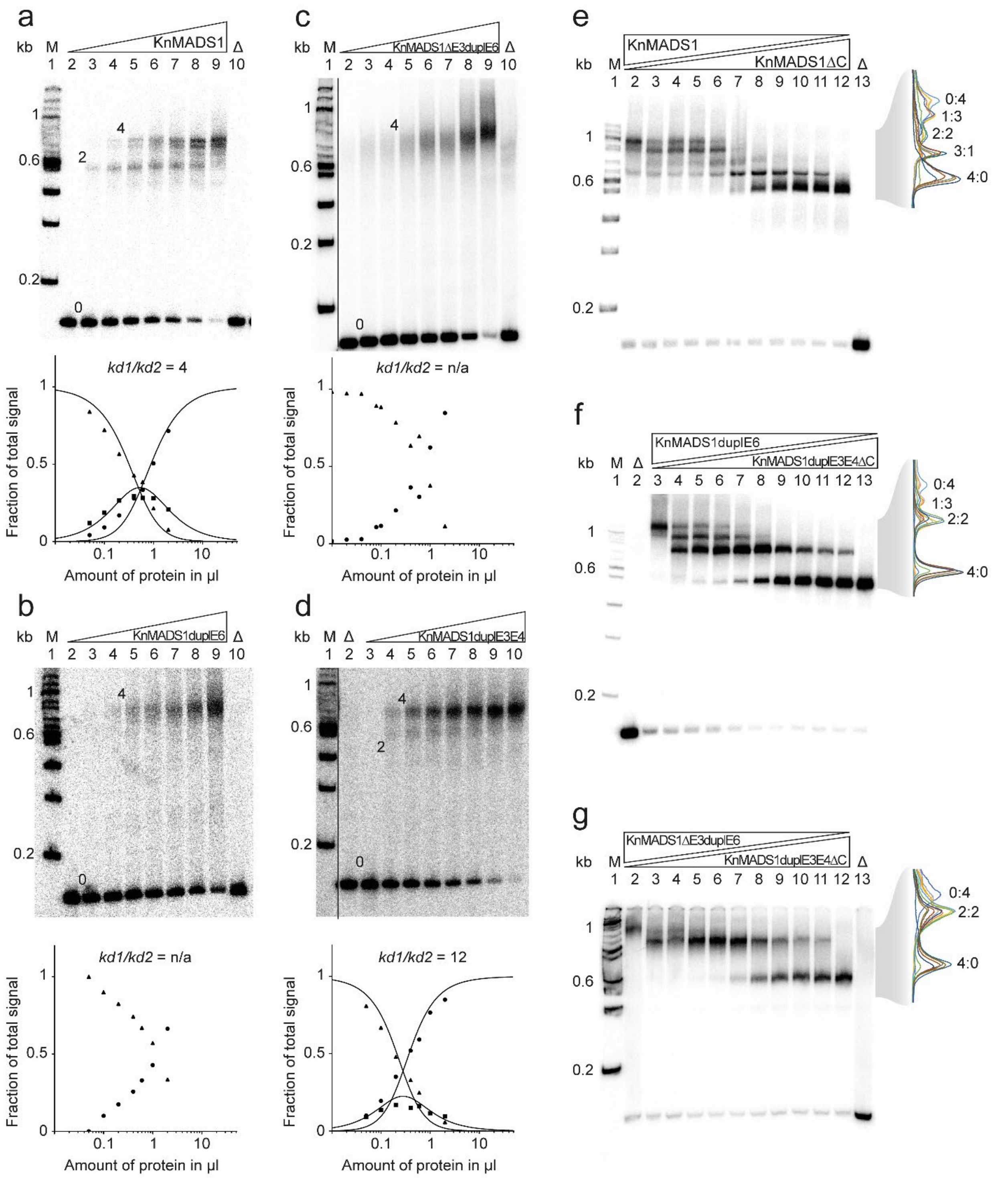


**Supplementary Figure 4: Original gel images of Figure 5.** See figure legend of Figure 5. For size comparison radioactively labelled 100 bp DNA ladder (NEB) was added ('M').

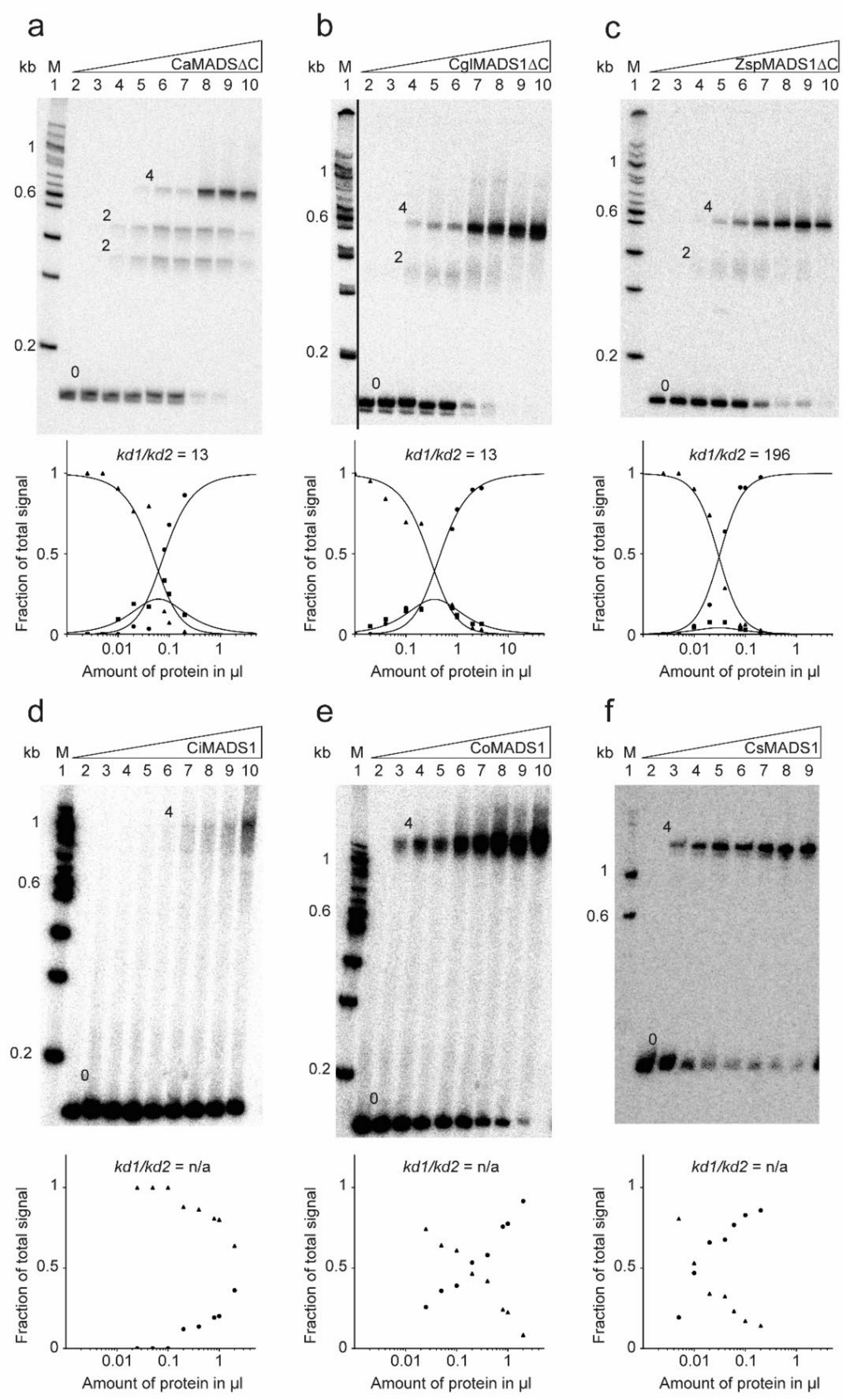


**Supplementary Figure 5: Original gel images of Figure 6.** See figure legend of Figure 6. For size comparison radioactively labelled 100 bp DNA ladder (NEB) was added ('M').
